# Identification of positive selection in genes is greatly improved by using experimentally informed site-specific models

**DOI:** 10.1101/037689

**Authors:** Jesse D. Bloom

## Abstract

Sites of positive selection are identified by comparing observed evolutionary patterns to those expected under a null model for evolution in the absence of such selection. For protein-coding genes, the most common null model is that nonsynonymous and synonymous mutations fix at equal rates; this unrealistic model has limited power to detect many interesting forms of selection. I describe a new approach that uses a null model based on high-throughput lab measurements of a gene's site-specific amino-acid preferences. This null model makes it possible to identify diversifying selection for amino-acid change and differential selection for mutations to amino acids that are unexpected given the measurements made in the lab. I show that this approach identifies sites of adaptive substitutions in four genes (lactamase, Gal4, influenza nucleoprotein, and influenza hemagglutinin) far better than a comparable method that simply compares the rates of nonsynonymous and synonymous substitutions. As rapid increases in biological data enable increasingly nuanced descriptions of the constraints on individual sites, approaches like the one here can improve our ability to identify many interesting forms of selection.

## Introduction

An important goal of biology is to identify genetic modifications that have led to interesting changes in phenotype. In the case of protein-coding genes, this means identifying mutations that were fixed by selection to alter properties such as the activity of enzymes or the antigenicity of viral proteins.

This goal is challenging because not all mutations that fix do so because they confer beneficial phenotypic effects that are selected by evolution. Sometimes mutations fix because they adaptively alter phenotype, but mutations also fix due to forces such as genetic drift or hitchhiking. Therefore, it is difficult to examine gene sequences and pinpoint specific mutations that have changed evolutionarily relevant phenotypes. As Zuckerkandl and Pauling (1965) noted a half-century ago:

> [Many] substitutions may lead to relatively little functional change, whereas at other times the replacement of one single amino acid residue by another may lead to a radical functional change… It is the type rather than number of amino acid substitutions that is decisive.

Unfortunately, Zuckerkandl and Pauling (1965) did not provide a prescription for determining the “type” of substitution that leads to phenotypic change, and such a prescription remains elusive decades later.

Because it is difficult to determine *a priori* which substitutions have altered relevant phenotypes, methods have been devised that compare homologous sequences to identify sites where mutations have been positively selected by evolution. The basic strategy is to formulate a null model for evolution, and then identify sites that have evolved in ways incompatible with this model. If the null model adequately describes evolution in the absence of selection for phenotypic change, then sites that deviate from the model are ones where mutations have been selected because they alter evolutionarily relevant phenotypes.

For protein-coding genes, the most widely used algorithms are built around the null model that nonsynonymous and synonymous mutations should fix at equal rates. These algorithms estimate the rates of fixation of nonsynonymous (*dN*) and synonymous (*dS*) mutations at each codon site *r* (Nielsen and Yang, 1998; Suzuki and Gojobori, 1999; Yang et al., 2000; Pond and Frost, 2005; Murrell et al., 2012). The ratio *dN*/*dS* at *r* is taken as a measure of selection. If the ratio is clearly > 1 then pressure for phenotypic change is favoring fixation of protein-altering nonsynonymous mutations, and the site is under diversifying selection. If the ratio is clearly < 1 then nonsynoymous mutations are being purged to prevent phenotypic change, and the site is under purifying selection.

Although *dN*/*dS* methods are tremendously useful (the leading software implementations HyPhy and PAML have each been cited thousands of times; Pond et al., 2005; Yang, 2007), their underlying null model is clearly oversimplified. A random amino-acid mutation completely inactivates the typical protein ≈40% of the time (Guo et al., 2004). So unsurprisingly, most genes have many sites with *dN*/*dS* < 1. This finding is often of little biological value, since researchers frequently already know that the gene they are studying is under some type of protein-level constraint. So detecting purifying selection as manifested by *dN*/*dS* < 1 often points more to the naivety of the null model than unexpected biology.

Perhaps more importantly, *dN*/*dS* methods also have limited power to identify sites that have fixed adaptive mutations. For instance, T-cells drive fixation of immune-escape mutations in influenza – but because the relevant sites are under strong constraint, *dN*/*dS* remains < 1 and the relative increase in nonsynonymous substitution rate is only apparent in comparison to homologs not subject to immune selection (Machkovech et al., 2015). Therefore, even positive selection for adaptive mutations can fail to elevate *dN*/*dS* > 1 at functionally constrained sites.

The limitations of a null model that assumes equal rates of fixation of nonsynonymous and synonymous mutations have become especially glaring in light of deep mutational scanning experiments. These experiments, which subject libraries of mutant genes to selection in the lab and query the fate of each mutation by deep sequencing (Fowler and Fields, 2014; Boucher et al., 2014), can measure the preference of each site in a protein for each amino acid (Bloom, 2015). A clear result is that sites vary wildly in their amino-acid preferences. Some sites are relatively unconstrained and prefer all amino acids roughly equally; for these sites, the null model of *dN*/*dS* methods is reasonable. But most sites strongly prefer one or a few amino acids; *dN*/*dS* methods do not offer a plausible null model for these sites.

As an example, Figure 1 shows the amino-acid preferences of five sites in TEM-1 ,0-lactamase as measured by the deep mutational scanning of Stiffler et al. (2015). Mutations at three of these sites confer antibiotic or inhibitor resistance in lactamases (Salverda et al., 2010). Inspection of Figure 1 shows that the two sites not implicated in resistance have evolved in ways that seem roughly compatible with their amino-acid preferences measured in the lab: site 201 tolerates many amino acids in the lab and is moderately variable in nature, while site 242 strongly prefers glycine in the lab and is conserved at that identity in nature. But the three sites involved in resistance have evolved in ways that seem to deviate from their amino-acid preferences measured in the lab: site 238 substitutes from the lab-preferred glycine to the less preferred serine, site 240 repeatedly substitutes to lysine despite not strongly preferring this amino acid in the lab, and site 244 substitutes from the lab-preferred arginine to several less preferred amino acids. So given the experimentally measured preferences, it is fairly apparent that the sites where mutations contribute to antibiotic resistance are evolving in ways that deviate from the preferences measured in the lab. But as Figure 1 shows, a *dN*/*dS* method fails to find any site with *dN*/*dS* > 1 at a false-discovery rate (FDR) of 0.05. As this example shows, a null model that fails to account for site-specific amino-acid preferences can overlook sites that fix adaptive mutations.

**Figure 1:**
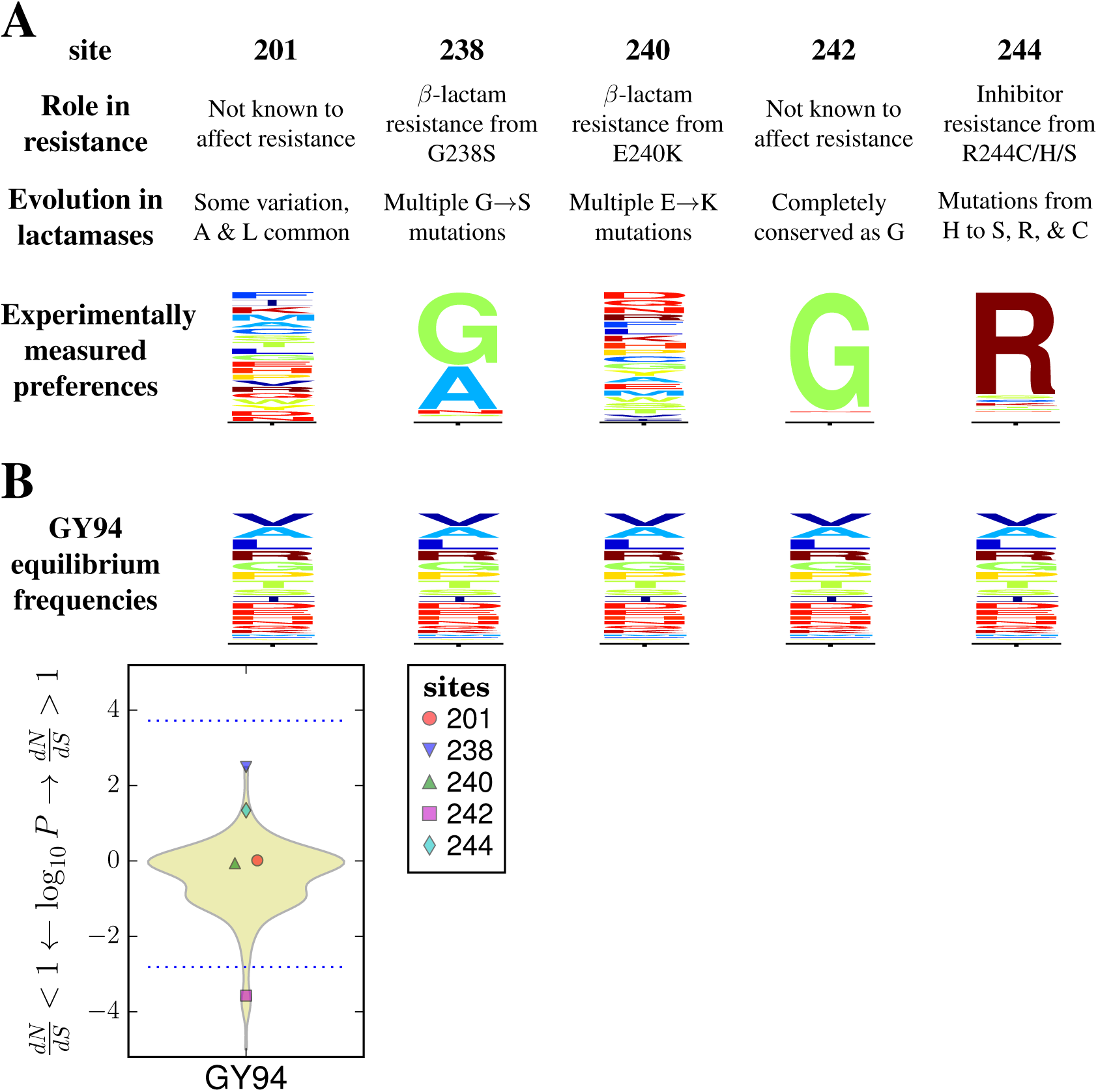
Different sites are expected to evolve differently, but *dN*/*dS* methods ignore this fact and so have limited power to detect positive selection. **(A)** The amino-acid preferences of five sites in TEM-1 *β*-lactamase as measured by Stiffler et al. (2015) (letter heights are proportional to amino-acid preferences). Three sites experience mutations that confer extended-spectrum antibiotic or inhibitor resistance (Salverda et al., 2010). The two sites not involved in resistance are evolving in a way that seems roughly compatible with their amino-acid preferences, while the three sites implicated in resistance are evolving in ways that clearly deviate from the preferences (for instance, site 238 mutates from highly preferred glycine to the very low preference amino-acid serine). **(B)** A standard *dN*/*dS* model (that of Goldman and Yang, 1994, abbreviated GY94) assumes all sites evolve under uniform constraints. When this model is used to fit a site-specific *dN*/*dS*, no sites are deemed under diversifying selection (*dN*/*dS* > 1) at a FDR of 0.05 for testing all sites, although the non-resistance site 242 is deemed under purifying selection (*dN*/*dS* < 1). The violin plot shows the distribution of P-values for sites having *dN*/*dS* > or < 1. All sites below the bottom dotted blue line are deemed to have *dN*/*dS* < 1 at and FDR of 0.05. No sites have *dN*/*dS* > 1 at this FDR, so the top dotted blue line indicate the *P*-value that would be needed for a site to have *dN*/*dS* > 1 at a significance level of 0.05 using a Bonferroni correction. This figure shows a handful of selected sites to illustrate the general principle, and does not represent a rigorous statistical analysis of the full gene. A full analysis of all sites and further details are later in the paper. See Supplementary figure 1 for subtleties about amino-acid preferences versus equilibrium frequencies.

Here I describe how the limitations of *dN*/*dS* methods illustrated in Figure 1 can be overcome. The approach that I describe defines selection relative to a null model established by experimentally measured site-specific amino-acid preferences, and so tests if sites are evolving differently in nature than expected from constraints measured in the lab. This more nuanced null model can be used to identify sites of *diversifying selection* for unusually rapid amino-acid change via a statistically principled extension to standard *dN*/*dS* methods. The more nuanced null model can also be used to heuristically identify sites of *differential selection* for unexpected amino acids. I apply the approach to four genes, and show that it greatly outperforms a *dN*/*dS* method at identifying sites of antibiotic-resistance and immune-escape mutations. As deep mutational scanning data become more widespread, approaches like the one here can enhance our ability to identify sites of biologically interesting selection.

## New Approaches

### An evolutionary null model informed by experimentally measured amino-acid preferences

To remedy the limitations of *dN*/*dS* methods illustrated in Figure 1, we formulate a description of how sites should evolve if selection in nature matches the constraints measured by deep mutational scanning in the lab. This description consists of a set of site-specific experimentally informed codon models (ExpCM). The ExpCM used here are similar but not identical to those in Bloom (2014a,b).

Deep mutational scanning experiments provide direct measurements of the preference *π*_*r*_,_*a*_ of each site *r* for each amino acid *a* (for details of how these preferences can be obtained from the experimental data, see Bloom, 2015). These preferences are normalized so Σ_*a*_ *π*_*r*_,_*a*_ = 1. We use the preferences to define an ExpCM for each site. As is typical for phylogenetic substitution models, each ExpCM is a reversible stochastic matrix giving the rates of substitution between codons. The rate *P*_*r*_,_*xy*_ from codon *x* to *y* at site *r* is written in mutation-selection form as

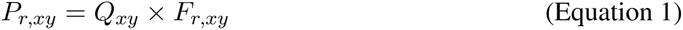

where *Q*_*xy*_ represents the rate of mutation from *x* to *y* and *F*_*r*_,_*xy*_ represents the selection on this mutation. The mutation terms are identical across sites, but the selection terms are site-specific.

The mutation terms *Q*_*xy*_ are given by a HKY85 model (Hasegawa et al., 1985), and consist of a transition-transversion ratio *κ* and four nucleotide parameters *ϕ*_*A*_, *ϕ*_*C*_, *ϕ*_*G*_, and *ϕ*_*T*_ that sum to one. These *ϕ* parameters give the expected nucleotide composition in the absence of selection on amino acids; the actual nucleotide frequencies are also influenced by the selection (for this reason, the *ϕ* terms cannot simply be equated with the empirical alignment frequencies as is sometimes done for conventional substitution models). The mutation term is:

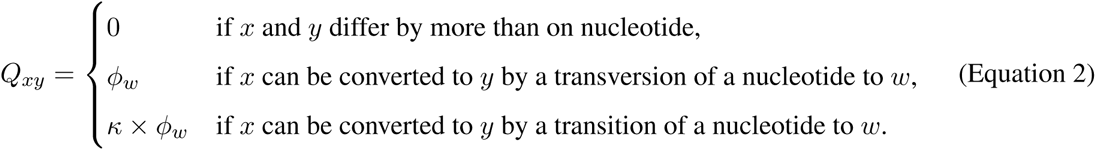

The site-specific amino-acid preferences *π*_*r*_,_*a*_ enter the model via the selection terms *F*_*r*_,_*xy*_. Let A (*x*) denote the amino acid encoded by codon *x*, let *β* be the stringency parameter described in Bloom (2014b), and let *ω* be a gene-wide relative rate of fixation of nonsynonymous to synonymous mutations after accounting for the amino-acid preferences. Then:

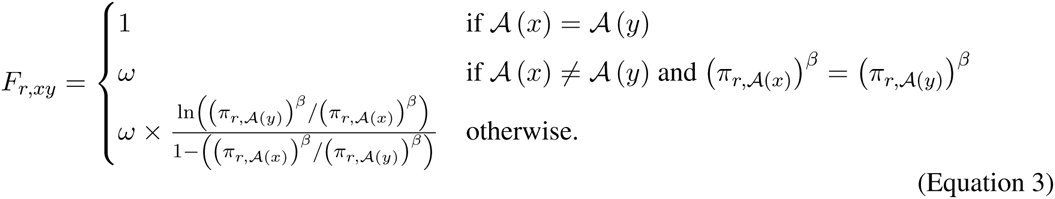

The functional form relating *F*_*r*_,_*xy*_ to *π*_*r*_,_*a*_ for nonsynonymous mutations is that derived by Halpern and Bruno (1998) under certain (probably unrealistic) assumptions about the evolutionary process and the relationship between the preferences and amino-acid fitnesses (see also McCandlish and Stoltzfus, 2014; Thorne et al., 2007; Yang and Nielsen, 2008). Relative to the equation of Halpern and Bruno (1998), Equation 3 removes terms related to mutation (these are captured by *Q*_*xy*_) and corrects a typographical error in the denominator. The stringency parameter *β* is > 1 if natural selection favors high-preference amino acids with greater stringency than the experiments used to measure *π*_*r*_,_*a*_, and is < 1 if it favors them with less stringency. Under the assumptions of Halpern and Bruno (1998), *β* is related to effective population size. Note that if *β* = 0, then the substitution model defined by Equation 1 reduces to a F1X4 version of the Goldman and Yang (1994) model. The ω parameter indicates if there is a retardation (*ω* < 1) or acceleration (*ω* > 1) in the rate of fixation of nonsynonymous mutations relative to synonymous mutations after accounting for the preferences. Bloom (2014b) shows that a model of the form defined by *P*_*r*,*xy*_ is reversible and has stationary state

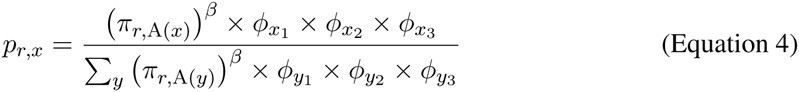

where *x*_1_, *x*_2_, and *x*_3_ are the nucleotides at positions 1, 2, and 3 of codon *x*.

The ExpCM can be used to calculate the likelihood of a phylogenetic tree and an alignment of genes using the algorithm of Felsenstein (1981), which implicitly assumes that sites evolve independently. The set of ExpCM for a given gene have six free parameters: *ω*, *β*, *κ*, and three of the *ϕ*’s. The *π*_*r*_,_*a*_ values are not free parameters, since they are specified *a priori* from experimental data. The values of the six free parameters are fit by maximum likelihood.

Overall, ExpCM describe how sites evolve if selection in nature is concordant with the amino-acid preferences measured in the lab. They therefore enable the visual comparison between the preferences and natural evolution in Figure 1 to be performed in a formal statistical manner.

### Identifying sites of diversifying selection

Having established a null model for how a gene should evolve if selection adheres to the constraints measured in the lab, we next want to identify sites that deviate from this model. Such sites are likely targets of additional selection. One such form of selection is *diversifying selection* for amino-acid change, as occurs at viral epitopes under continual pressure to escape newly generated immunity.

To detect diversifying selection, we use an approach analogous the fixed effects likelihood (FEL) method (Pond and Frost, 2005; Massingham and Goldman, 2005; Suzuki, 2004). After fixing the tree and model parameters to their maximum likelihood values for the entire sequence, for each site *r* we fit a synonymous rate *μ*_*r*_ and a parameter *ω*_*r*_ corresponding to the nonsynonymous rate relative to the synonymous rate by replacing Equation 3 with

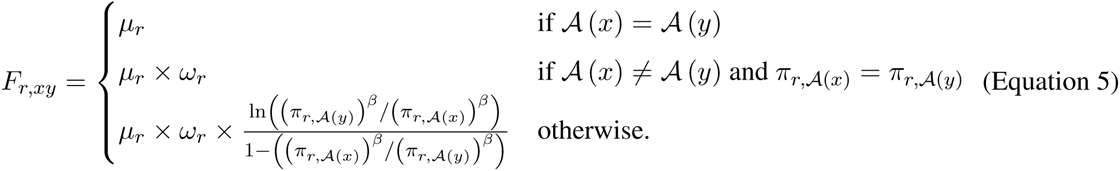

and optimizing with respect *μ*_*r*_ and *ω*_*r*_. The reason that we fit *μ*_*r*_ as well as *ω*_*r*_ is to accommodate synonymous rate variation among sites; this can be important for the reasons described by Pond and Muse (2005). The null hypothesis is that *ω*_*r*_ = 1. Following Pond and Frost (2005), we compute a P-value for rejecting this null hypothesis by using a 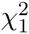 test to compare the likelihood when fitting both *μ*_*r*_ and *ω*_*r*_ to that when fitting only *μ*_*r*_ and fixing *ω*_*r*_ = 1. The key statistic is not *ω*_*r*_ itself, but rather the P-value for rejecting the null hypothesis in favor of *ω*_*r*_ > 1 or *ω*_*r*_ < 1. The former case implies diversifying selection, while the latter case indicates a selective constraint on amino-acid change that is not adequately captured by the preferences. To account for the fact that a different test is performed for each site, we control the FDR using the procedure of Benjamini and Hochberg (1995). As demonstrated below, this approach has excellent power to pinpoint sites like 238 and 244 in Figure 1, which fix multiple amino-acid mutations despite being under strong functional constraint.

### Identifying sites of differential selection

Some interesting forms of selection do not cause sites to change repeatedly, but rather lead them to substitute to amino acids that are unexpected given the amino-acid preferences measured in the lab. Such sites are under *differential selection* to fix mutations different from those expected if selection in nature parallels that in the lab.

To detect differential selection, we compare the preferences measured in the lab to those that optimally describe evolution in nature. Denote the preferences that optimally describe evolution in nature as *π̂*_*r*,*a*_ with Σ_*a*_*π̂*_*r*,*a*_ = 1. Denote the differential preference Δ*π*_*r*_,_*a*_ for amino-acid *a* at site *r* as the difference between *π̂*_*r*,*a*_ and the experimentally measured preferences rescaled by the stringency parameter: 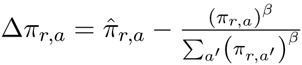. If we redefine Equation 3 by replacing (*π*_*r*_,_*a*_)^*β*^ with *π̂*_*r*_,_*a*_ as in

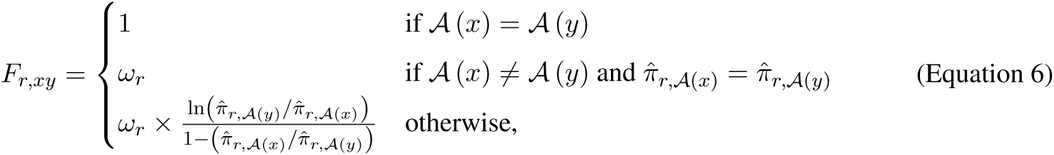

then we can determine the preferences that optimally describe natural evolution by optimizing with respect to *π̂*_*r*_,_*a*_ after fixing the tree and model parameters to their maximum likelihood values for the entire sequence. However, unconstrained optimization of Equation 6 will overfit the data. We therefore instead optimize the product of Equation 6 and an equation that regularizes the Δ*π*_*r*_,_*a*_ values by biasing them towards zero:

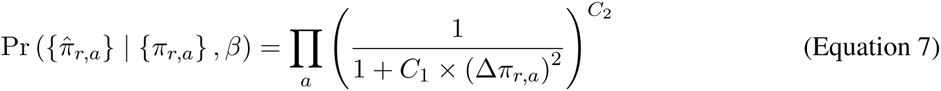

where *C*_1_ and C_2_ determine how strongly *π̂*_*r*_,_*a*_ is biased towards the experimentally measured preferences. Here I use *C*_1_ = 150 and *C*_2_ = 0.5; Equation 7 is illustrated in Supplementary figure 2. Note that while the underlying rationale for regularizing the Δ*π*_*r*_,_*a*_ values is clear, the regularization implemented by Equation 7 was chosen heuristically (a more statistically principled method for regularizing the Δ*π*_*r*_,_*a*_ values is an important area for future work).

A differential preference of Δ*π*_*r*_,_*a*_ > 0 implies that natural evolution favors amino-acid *a* at site *r* more than expected, whereas Δ*π*_*r*_,_*a*_ < 0 implies that evolution disfavors this amino acid. The total differential selection at *r* is quantified as half the absolute sum of the differential preferences, 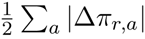; this quantity ranges from zero to one. As demonstrated below, this approach has excellent power to pinpoint sites like 238 and 240 in Figure 1, which fix mutations to unexpected amino acids. However, I emphasize that the test for differential selection is currently heuristic, and does not incorporate formal statistical significance testing.

## Results

### Choice of four genes to test approaches to identify sites of selection

To test the approaches for detecting selection described above, I selected four genes: the DNA-binding domain of yeast Gal4, *β*-lactamase, the nucleoprotein (NP) of human influenza, and the hemagglutinin (HA) of human seasonal H1N1 influenza. Previous deep mutational scanning studies have measured the effects of all mutations to these genes (Kitzman et al., 2015; Stiffler et al., 2015; Doud et al., 2015; Thyagarajan and Bloom, 2014), enabling calculation of their site-specific amino-acid preferences. For each gene, I assembled an alignment of homologs for evolutionary analysis (Table 1).

**Table 1:**
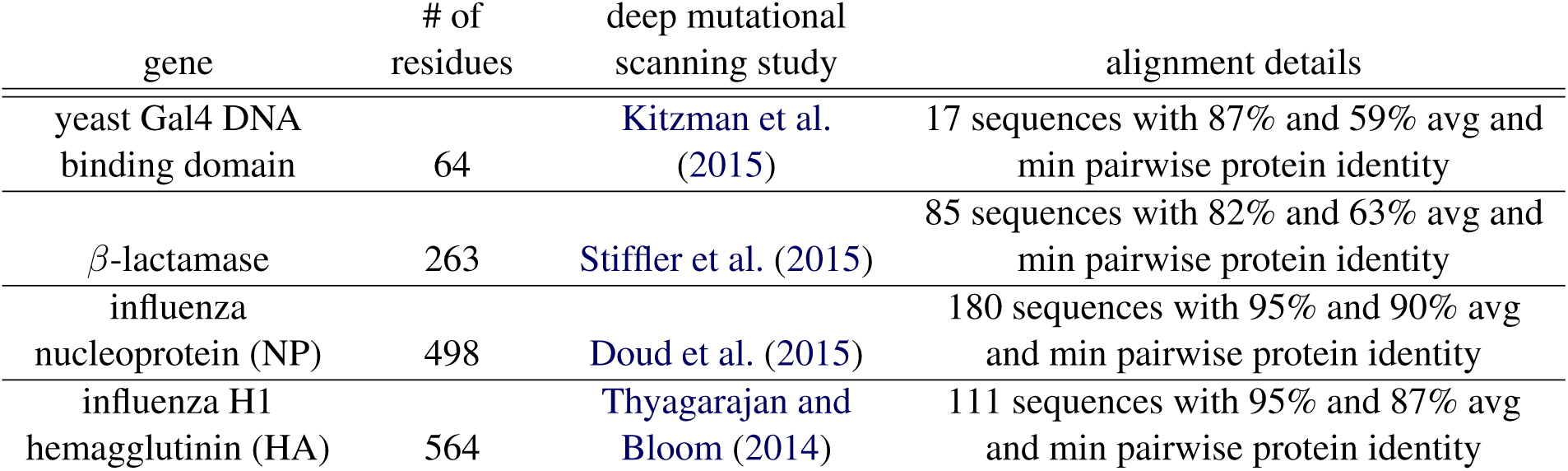
The four genes analyzed in this study.

| gene | # of residues | deep mutational scanning study | alignment details |
| --- | --- | --- | --- |
| yeast Gal4 DNA binding domain | 64 | Kitzman et al. (2015) | 17 sequences with 87% and 59% avg and min pairwise protein identity |
| β-lactamase | 263 | Stiffler et al. (2015) | 85 sequences with 82% and 63% avg and min pairwise protein identity |
| influenza nucleoprotein (NP) | 498 | Doud et al. (2015) | 180 sequences with 95% and 90% avg and min pairwise protein identity |
| influenza H1 hemagglutinin (HA) | 564 | Thyagarajan and Bloom (2014) | 111 sequences with 95% and 87% avg and min pairwise protein identity |

A great deal is known about the pressures that have shaped the evolution of all four genes. Gal4 performs a function that is conserved among homologs from widely diverged species, and does not appear to be changing phenotypically (Johnston, 1987; Traven et al., 2006). However, the other three genes are undergoing adaptive evolution: lactamases evolve resistance to new antibiotics and inhibitors (Du Bois et al., 1995; Salverda et al., 2010), while NP and HA evolve to escape the immune response in humans (Voeten et al., 2000; Machkovech et al., 2015; Yewdell et al., 1979; Caton et al., 1982). These genes therefore provide an excellent test case. Gal4 is a “negative control”: no sites in this gene should be identified as under selection to fix adaptive mutations. But an effective approach for identifying positive selection should pinpoint the sites of drug-resistance and immune-escape mutations in the other three genes.

### Experimentally informed site-specific models are vastly better descriptors of evolution

Our basic assumption is that site-specific ExpCM are a better null model for evolution than the nonsite-specific models used by *dN*/*dS* methods. Prior work has shown that experimentally informed site-specific models similar to the ExpCM defined here greatly outperform non-site-specific models (Bloom, 2014a,b; Thyagarajan and Bloom, 2014; Doud et al., 2015). To confirm this result for the ExpCM and genes here, I compared the ExpCM to the model of Goldman and Yang (1994) (denoted as GY94), which is commonly used by *dN*/*dS* methods. I used F3X4 equilibrium frequencies for GY94, with the nine F3X4 parameters estimated by maximum likelihood. These equilibrium frequencies are *not* site-specific; this is the major difference between GY94 and ExpCM (Figure 1).

To compare the models and perform the other analyses in this paper, I developed the software package `phydms` (**<u>phy</u>**logenetics informed by **<u>d</u>**eep **<u>m</u>**utational **<u>s</u>**canning; https://github.com/jbloom/phydms). This software interfaces with and extends `Bio`++ (Dutheil et al., 2006; Guéguen et al., 2013) to enable analyses with both ExpCM and GY94 models.

I used phydms to infer a maximum-likelihood phylogenetic tree for each gene using GY94 with a single gene-wide *dN*/*dS* ratio (the M0 model of Yang et al. (2000)). After fixing the tree topology to that estimated using GY94 M0, I re-optimized the branch lengths and model parameters by maximum likelihood for four additional models. The first is GY94 M3 (Yang et al., 2000), in which the likelihood for each site is a linear combination of those under three different *dN*/*dS* values, with these values and their weights shared across the whole alignment and optimized by maximum likelihood. The second is ExpCM. The third is ExpCM with the amino-acid preferences averaged across sites – this averaging makes the model non-site-specific, but captures any gene-wide trends in the deep mutational scanning data. The final is ExpCM with the amino-acid preferences randomized among sites – this model is still site-specific, but the site-specific parameters are no longer associated with the actual site for which they were measured.

I compared these models using Akaike Information Criteria (AIC) (Posada and Buckley, 2004), which measures model fit penalized by the number of free parameters. Table 2 shows that ExpCM describe the evolution of all four genes far better than any other model. The huge superiority of ExpCM over the GY94 models is because ExpCM capture site-specific evolutionary constraints, as demonstrated by the fact that ExpCM in which preferences are averaged across sites are comparable to GY94. The failure of the averaged ExpCM to offer consistent improvement over GY94 is because most specificity in the substitution process occurs at the level of sites, not the level of genes – so there is little to be gained from a gene-specific model that treats all sites the same. The poor performance of the randomized ExpCM is because a site-specific model only helps if the experimentally measured preferences are assigned to the correct sites. Indeed, Table 2 shows that randomly assigned site-specific preferences are so detrimental that they are nearly completely flattened by fitting a stringency parameter *β* that is close to zero, effectively making the randomized ExpCM non-site-specific. Overall, Table 2 confirms previous work (Bloom, 2014a,b; Thyagarajan and Bloom, 2014; Doud et al., 2015) showing that experimentally informed site-specific models provide vastly improved descriptions of evolution.

**Table 2:** The site-specific ExpCM describe the evolution of all four genes vastly better than GY94 or ExpCM in which the amino-acid preferences have been averaged or randomized across sites.

**Gal4**
| model | ΔAIC | log likelihood | # free parameters: values of selection parameters |
| --- | --- | --- | --- |
| ExpCM | 0 | -1048 | 6: $\beta = 0.82, \omega = 0.13$ |
| GY94 M3 | 129 | -1103 | 15: $\omega_1 = 0.01, \omega_2 = 0.11, \omega_3 = 0.49, p_1 = 0.50, p_2 = 0.29$ |
| GY94 M0 | 192 | -1139 | 11: $\omega = 0.06$ |
| averaged ExpCM | 196 | -1146 | 6: $\beta = 1.07, \omega = 0.06$ |
| randomized ExpCM | 206 | -1151 | 6: $\beta = 0.10, \omega = 0.07$ |

**lactamase**
| model | ΔAIC | log likelihood | # free parameters: values of selection parameters |
| --- | --- | --- | --- |
| ExpCM | 0 | -3421 | 6: $\beta = 1.01, \omega = 1.02$ |
| GY94 M3 | 564 | -3694 | 15: $\omega_1 = 0.07, \omega_2 = 0.55, \omega_3 = 6.24, p_1 = 0.69, p_2 = 0.17$ |
| GY94 M0 | 765 | -3798 | 11: $\omega = 0.34$ |
| averaged ExpCM | 766 | -3804 | 6: $\beta = 0.77, \omega = 0.35$ |
| randomized ExpCM | 790 | -3816 | 6: $\beta = 0.10, \omega = 0.34$ |

**NP**
| model | ΔAIC | log likelihood | # free parameters: values of selection parameters |
| --- | --- | --- | --- |
| ExpCM | 0 | -8624 | 6: $\beta = 2.43, \omega = 0.61$ |
| GY94 M3 | 2175 | -9703 | 15: $\omega_1 = 0.00, \omega_2 = 0.16, \omega_3 = 1.31, p_1 = 0.59, p_2 = 0.24$ |
| averaged ExpCM | 2584 | -9916 | 6: $\beta = 0.43, \omega = 0.11$ |
| randomized ExpCM | 2593 | -9921 | 6: $\beta = 0.10, \omega = 0.11$ |
| GY94 M0 | 2613 | -9926 | 11: $\omega = 0.11$ |

**HA**
| model | ΔAIC | log likelihood | # free parameters: values of selection parameters |
| --- | --- | --- | --- |
| ExpCM | 0 | -7461 | 6: $\beta = 1.61, \omega = 0.60$ |
| GY94 M3 | 1782 | -8343 | 15: $\omega_1 = 0.02, \omega_2 = 4.26, \omega_3 = 4.94, p_1 = 0.59, p_2 = 0.25$ |
| averaged ExpCM | 2137 | -8530 | 6: $\beta = 0.42, \omega = 0.23$ |
| randomized ExpCM | 2157 | -8539 | 6: $\beta = 0.10, \omega = 0.23$ |
| GY94 M0 | 2176 | -8544 | 11: $\omega = 0.22$ |

Another informative comparison is between the *dN*/*dS* of GY94 and the *ω* of ExpCM. ExpCM can represent protein-level constraint either via the site-specific amino-acid preferences or by shrinking *ω* to < 1. In contrast, GY94 can only represent constraint by shrinking *dN*/*dS* even if the actual selection is for preferred amino acids at each site rather than against amino-acid change *per se* (Spielman and Wilke, 2015a). Table 2 shows that the ExpCM *ω* is always greater than the GY94 *dN*/*dS*. This effect is most striking for lactamase: while GY94 suggests selection against amino-acid change *per se* by fitting *dN*/*dS* = 0.3, ExpCM indicate that this selection is actually accounted for by the site-specific amino-acid preferences by fitting *ω* = 1. For the other three genes, the ExpCM *ω* is < 1 indicating that the site-specific amino-acid preferences don’t capture all constraints, but the ExpCM *ω* is still always substantially greater than the GY94 *dN*/*dS*.

The ExpCM stringency parameter *β* also provides useful information. Recall that *β* > 1 means that natural evolution selects for preferred amino acids with greater stringency than the deep mutational scanning. Table 2 shows that for both influenza genes (NP and HA), the stringency of natural evolution exceeds that of the deep mutational scanning, indicating that the experiments of Doud et al. (2015) and Thyagarajan and Bloom (2014) were not as rigorous as selection in nature. For lactamase, the stringency of natural evolution is approximately equal to that of the deep mutational scanning, providing a second indication (along with the fitting of *ω* ≈ 1) that the experiments of Stiffler et al. (2015) did an excellent job of capturing the constraints on lactamases in nature. Only for Gal4 is *β* < 1: either the selections of Kitzman et al. (2015) were more stringent than natural selection, or the measured preferences are not completely representative of those in nature and so is *β* fit to < 1 to somewhat flatten these preferences.

The stringency-rescaled amino-acid preferences are in Figure 2, Supplementary figure 3, Supplementary figure 4, and Supplementary figure 5. These figures reveal remarkable variation in constraint among sites, explaining why ExpCM better describe evolution than non-site-specific models. Overall, the results in this section verify that ExpCM offer a better evolutionary null model, and so motivate their use in identifying diversifying and differential selection.

**Figure 2:**
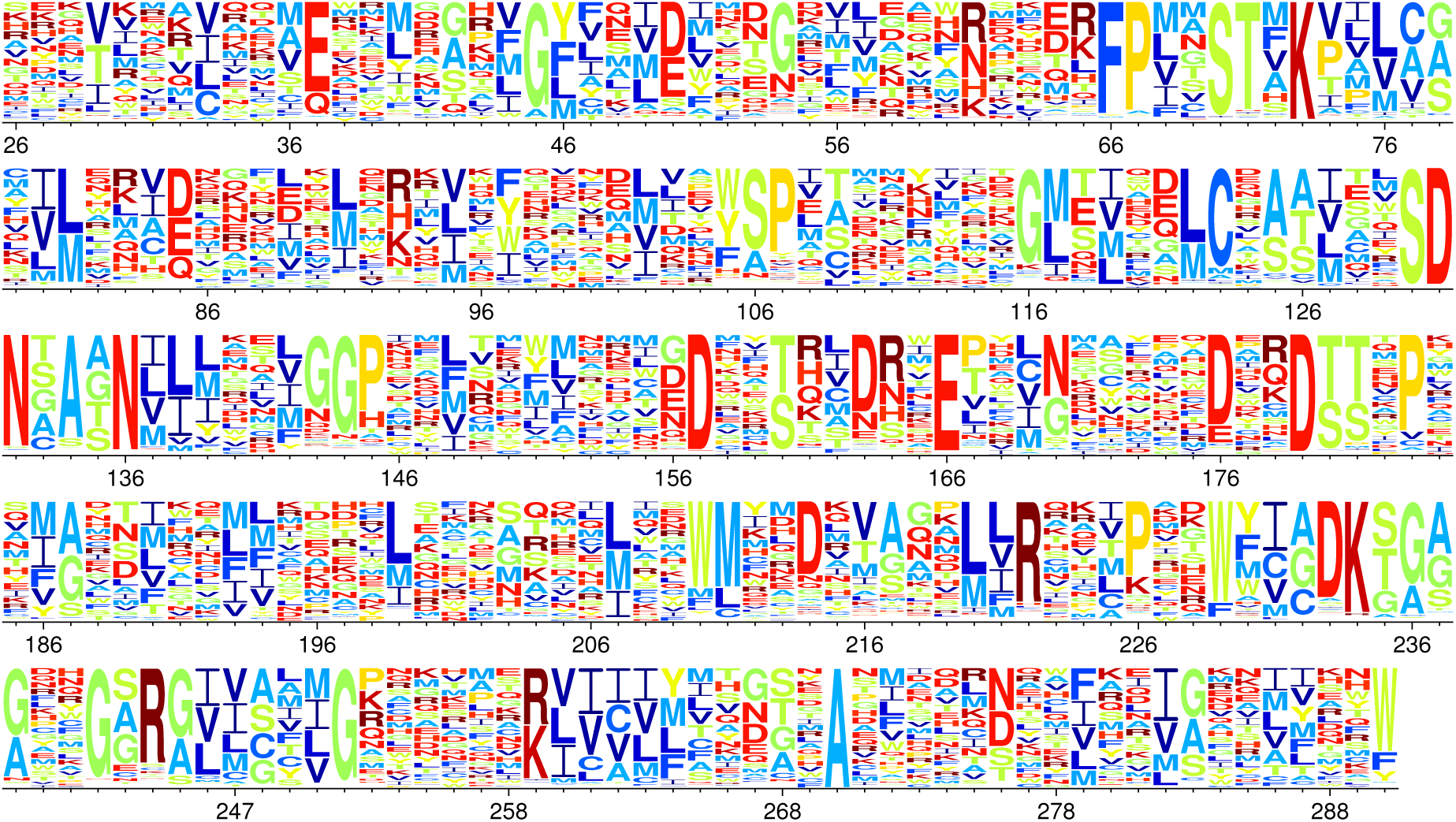
Site-specific amino-acid preferences for lactamase. The height of each letter is proportional to the preference for that amino acid at that site, and letters are colored by amino-acid hydrophobicity. These are the preferences experimentally measured by Stiffler et al. (2015) for TEM-1 *β*-lactamase under selection with 2.5 mg/ml ampicillin, re-scaled by the stringency parameter *β* = 1.01 from Table 2. The re-scaling is done so that if the experimentally measured preference for amino-acid *a* at site *r* is *π*_*r*,*a*_, then the rescaled preference is proportional to (*π*_*r*_,_*a*_)^*β*^. The lactamase sequence is numbered using the scheme of Ambler et al. (1991), meaning that residue numbers 239 and 253 are skipped. Comparable data for Gal4, NP, and HA are shown in Supplementary figure 3, Supplementary figure 4, and Supplementary figure 5, respectively.

### Experimentally informed site-specific models better detect diversifying selection

I used the ExpCM to identify sites of diversifying selection for amino-acid change. This was done by using phydms to fit *ω*_*r*_ and a synonymous rate for each site *r* via Equation 5, fixing all other parameters at their optimized values. To compare to a standard *dN*/*dS* method, I also fit a *dN*/*dS* ratio and synonymous rate for each site using GY94 with all other parameters fixed to the values optimized under GY94 M3 (equivalent to the fixed effects likelihood or FEL method as implemented by Pond and Frost (2005)).

Figure 3A shows that ExpCM have much greater power to identify diversifying selection than the GY94 *dN*/*dS* method. For Gal4, GY94 finds many sites with *dN*/*dS* < 1, but no sites with *dN*/*dS* > 1 at an FDR of 0.05. As discussed in the Introduction, identifying sites with *dN*/*dS* < 1 points to the naivety of the GY94 null model rather than unexpected biology, since any reasonable researcher would have already expected Gal4’s protein sequence to be under evolutionary constraint. The more plausible ExpCM null model finds that all sites in Gal4 are evolving as expected from the measurements in the lab (for no sites does it reject the null hypothesis *ω*_*r*_ = 1). For the other three genes, GY94 again finds that there are many sites with *dN*/*dS* < 1 while failing to identify any sites with *dN*/*dS* > 1 at an FDR of 0.05 – despite the fact that there is clear evidence that all three genes fix drug-resistance or immune-escape mutations. In contrast, the more realistic ExpCM find sites of diversifying selection for all three genes: there are three sites with *ω*_*r*_ > 1 in lactamase, four in NP, and two in HA.

**Figure 3:**
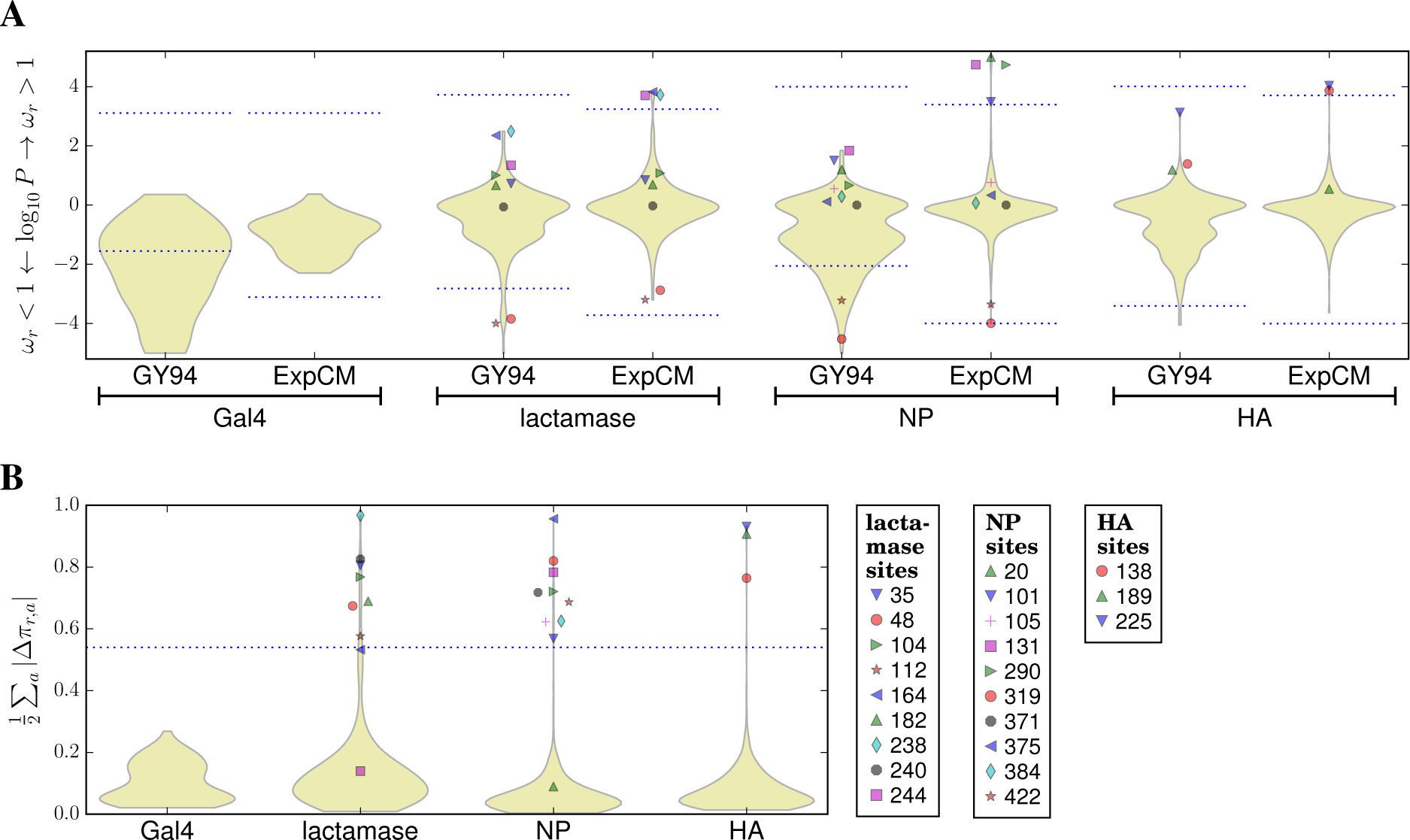
The experimentally informed models (ExpCM) identify many sites of diversifying or differential selection that are missed by a standard *dN*/*dS* analysis (GY94). **(A)** The violin plots show the distribution of *P*-values that a site is under diversifying selection for (positive numbers) or against (negative numbers) amino-acid change (*ω*_*r*_ indicates both the ExpCM parameter in Equation 5 and the GY94 *dN*/*dS* ratio). The portion of the distribution above / below the dotted blue lines contains all sites for which there is support for rejecting the null hypothesis *ω*_*r*_ = 1 at a FDR of 0.05. When there are no sites with support at this FDR, the dotted blue lines indicate the P-value that would be needed for a site to have *ω*_*r*_ > 1 or < 1 at a significance level of 0.05 using a Bonferroni correction. The *dN*/*dS* method identifies many sites of purifying selection, but fails to find any sites of selection for amino-acid change. The ExpCM model already accounts for basic functional constraints and so doesn’t identify any sites with *ω*_*r*_ < 1, but does identify sites of diversifying selection in all genes except Gal4 (which is not thought to evolve under pressure for phenotypic change). **(B)** The violin plots shown the distribution of differential selection at each site inferred with the ExpCM. Since Gal4 is not under selection for phenotypic change, I defined a heuristic threshold at 2-times the Gal4 maximum value of 0.27. At this threshold, sites of differential selection are identified for all three other genes. The legend labels all sites under diversifying or differential selection. This analysis was performed using `phydms`; Supplementary figure 6 shows that similar results are obtained if the *dN*/*dS* analysis is instead performed using `HyPhy` (Pond et al., 2005).

To statistically validate the ExpCM approach for identifying diversifying selection, I used pyvolve (Spielman and Wilke, 2015b) to simulate alignments of NP under ExpCM informed by the experimentally measured preferences and using the tree inferred for the actual NP sequences. In each simulation, I randomly selected five sites and placed them under diversifying with *ω*_*r*_ values ranging from 5 to 30. I then analyzed the simulated alignments for diversifying using the ExpCM and the FEL-like GY94 *dN*/*dS* method. As shown in Supplementary figure 7, ExpCM consistently outperformed GY94 at identifying the simulated sites of diversifying selection. Supplementary figure 7 also shows that the Benjamini and Hochberg (1995) procedure effectively controlled the false discovery rate. These simulations demonstrate the statistical soundness of the ExpCM approach for identifying diversifying selection.

Both the FEL-like GY94 *dN*/*dS* method and the ExpCM used for the analysis in Figure 3A test for diversifying selection across the phylogeny. But in many cases, diversifying selection is episodic. Therefore, *dN*/*dS* methods have been extended to identify sites under diversifying selection in only some lineages (Yang and Nielsen, 2002; Guindon et al., 2004; Yang and Dos Reis, 2011; Murrell et al., 2012). I used one of these methods, `MEME` (Murrell et al., 2012), to test for episodic diversifying selection. Supplementary figure 8 shows that `MEME` identifies one site of diversifying selection each in lactamase and NP, and no sites in HA or Gal4. This makes `MEME` more powerful than the FEL-like GY94 method but still less powerful than ExpCM. However, `MEME` and ExpCM outperform the FEL-like GY94 method for orthogonal reasons: `MEME` is superior because it can identify episodic selection, whereas ExpCM are superior because they account for functional constraints on individual sites. In principle, it should be possible to merge ExpCM with methods to identify episodic diversifying selection.

A variety of other *dN*/*dS* methods have also been developed. The most prominent other class includes so-called “random effects” methods that use an empirical Bayesian approach to share information about the distribution of *dN*/*dS* across sites (Nielsen and Yang, 1998; Yang et al., 2005; Huelsenbeck et al., 2006; Murrell et al., 2013). The relative pros and cons of “random effects” methods versus the “fixed effects” methods used in this paper remain an area of active discussion (Pond and Frost, 2005; Echave et al., 2016). It is beyond the scope of the current study to compare these two classes of methods. Here I simply note that as with the test for episodic selection described in the previous paragraph, ExpCM substitution models could in principle also be incorporated into the “random effects” framework, since the essential differences between “random effects” and “fixed effects” methods are due to how parameters are handled rather than the substitution model itself.

Overall, the results in this section show that ExpCM are better at identifying diversifying selection than several standard *dN*/*dS* methods. The reason for this superiority is that the ExpCM account for variation in the inherent constraints on different sites, and so have greater power to recognize when a functionally constrained site is changing more rapidly than expected.

### Experimentally informed site-specific models enable detection of differential selection

ExpCM also enable identification of differential selection for unexpected amino acids. I used phydms to estimate the differential preference Δ*π*_*r*_,_*a*_ of each site *r* for each amino-acid *a* by optimizing the product of Equation 6 and Equation 7 after fixing all other parameters. The differential selection at each site *r* was quantified as 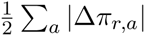, which can range from zero to one.

Figure 3B shows the distribution of site-specific differential selection. As expected, no sites in Gal4 are under strong differential selection. But for each of the other genes, a small subset of sites are under strong differential selection. I heuristically classified differential selection as “significant” if it exceeded 2-times the maximum value for Gal4. At this threshold, there are seven sites of differential selection in lactamase, nine in NP, and three in HA. So overall, Figure 3B suggests that most sites are evolving as expected in all four genes, but a small fraction of sites are under differential selection in lactamase, NP, and HA due to their roles in drug resistance or immune escape. This result is concordant from what we expect given biological knowledge about the selection pressures on these genes. Note that similarly reasonable results are *not* obtained using the non-phylogenetic Kullback-Leibler divergence to measure differences between amino-acid frequencies in nature and the experimentally measured aminoacid preferences (Supplementary figure 9). This fact emphasizes the importance of examining evidence for diversifying selection in a phylogenetic framework rather than simply treating natural sequences as an unbiased statistical ensemble.

A more detailed portrayal of the diversifying selection at each site is in Figure 4, Supplementary figure 10, Supplementary figure 11, and Supplementary figure 12. For each site, these images display the evidence for diversifying selection, the strength of differential selection, and the differential preference for each amino acid at sites under non-negligible differential selection.

**Figure 4:**
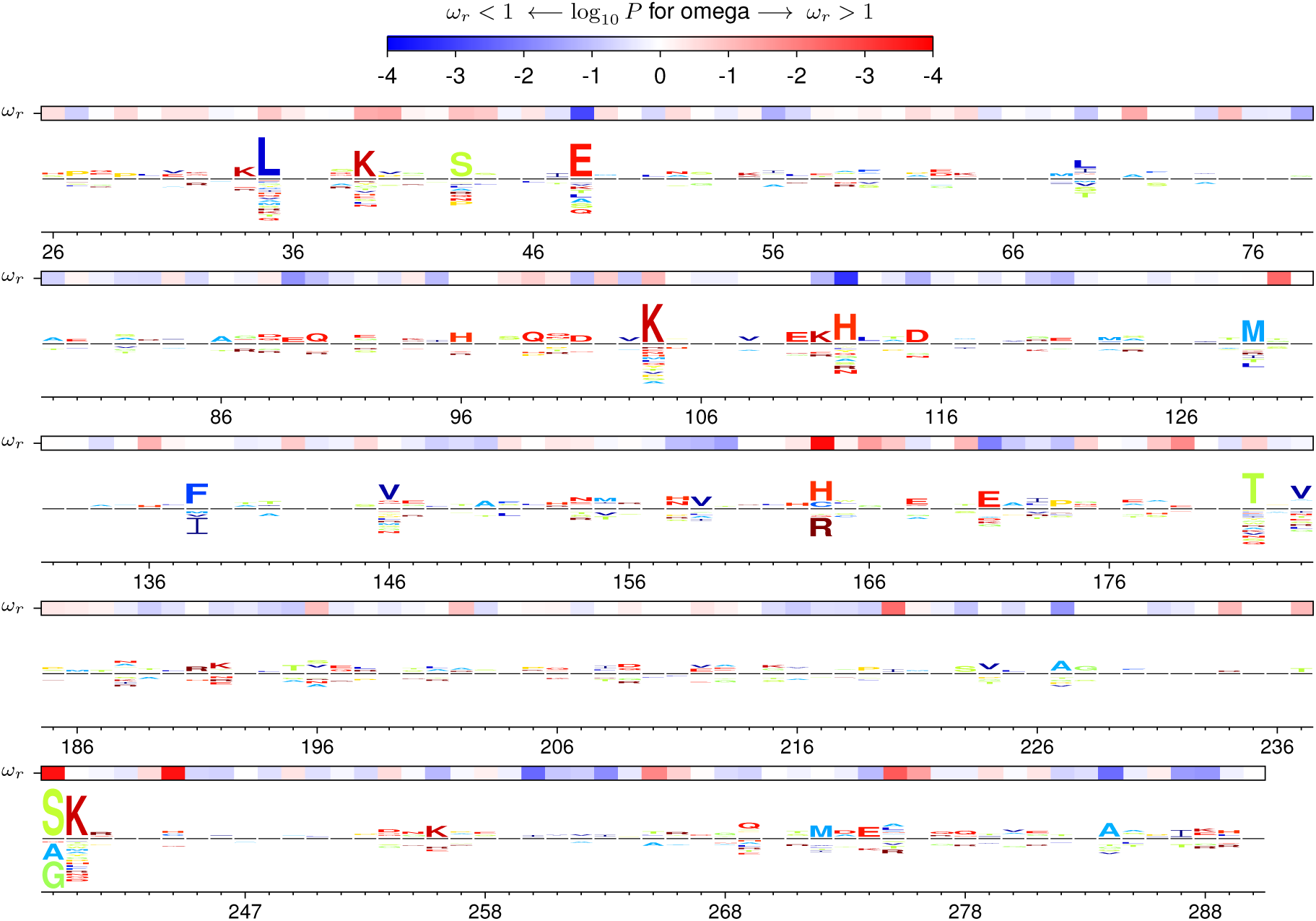
Site-specific selection on lactamase inferred with experimentally informed models. The height of each letter above/below the black center line is proportional to the differential selection for/against that amino acid at that site relative to what is expected from the amino-acid preferences in Figure 2. The overlay bar shows the evidence for diversifying selection at each site, which is manifested by strong evidence for a ratio *ω* of nonsynonymous to synonymous substitution rates that is higher (red) or lower (blue) than expected from the amino-acid preferences. The lactamase sequence is numbered using the scheme of Ambler et al. (1991), meaning that residue numbers 239 and 253 are skipped. Comparable data for Gal4, NP, and HA are shown in Supplementary figure 10, Supplementary figure 11, and Supplementary figure 12, respectively.

There are sites in lactamase, NP, and HA that are under both diversifying and differential selection, but there are also sites that are only under one of these forms of selection (Figure 3). These findings make sense: often, pressure for amino-acid change will drive multiple substitutions to non-preferred amino-acid identities, leaving traces of both types of selection. But sometimes, a relatively unconstrained site substitutes to a variety of different amino acids, leading to diversifying but not differential selection. In other cases, a site fixes just one or a few substitutions to a non-preferred amino acid that confers some enduring phenotypic benefit, leading to differential but not diversifying selection.

### The identified sites of selection are consistent with existing biological knowledge

The ExpCM identified sites of differential and diversifying selection in all three genes that are undergoing adaptive evolution (lactamase, NP, and HA), while GY94 identified no sites with *dN*/*dS* < 1 in any of the genes. But before concluding that this result indicates the superiority of the ExpCM, we must answer the following question: are the identified sites actually the locations of substitutions that have altered evolutionarily relevant phenotypes? To answer this question, I examined the literature on drug resistance in lactamases and immune escape by NP and HA (Table 3).

**Table 3:** Most sites of selection identified using the ExpCM are positions where mutations are known to affect biologically relevant drug resistance or immune escape phenotypes.

| gene | site | evidence for an effect on a biologically relevant phenotype |
| --- | --- | --- |
| lactamase | 35 | No evidence implicating this site in resistance (Salverda et al., 2010) |
| lactamase | 48 | No evidence implicating this site in resistance (Salverda et al., 2010) |
| lactamase | 104 | E104K involved in β-lactam resistance (Salverda et al., 2010) |
| lactamase | 112 | No evidence implicating this site in resistance (Salverda et al., 2010) |
| lactamase | 164 | R164C, R164H, and R164S involved in β-lactam resistance (Salverda et al., 2010) |
| lactamase | 182 | M182T potentiates resistance (Salverda et al., 2010) |
| lactamase | 238 | G238S involved in β-lactam resistance (Salverda et al., 2010) |
| lactamase | 240 | E240K involved in β-lactam resistance (Salverda et al., 2010) |
| lactamase | 244 | R244C, R244H, and R244S involved in inhibitor resistance (Salverda et al., 2010) |
| NP | 20 | Forms part of a T-cell epitope (Alexander et al., 2010) |
| NP | 101 | Forms part of an antibody epitope (Varich et al., 2011) |
| NP | 105 | Forms part of a T-cell epitope (Berkhoff et al., 2007) |
| NP | 131 | No reports of this site as a target of immune selection |
| NP | 290 | Forms part of an antibody epitope (Varich and Kaverin, 2004) |
| NP | 319 | No reports of this site as a target of immune selection |
| NP | 371 | Forms part of an antibody epitope (Miyoshi-Akiyama et al., 2012; Varich et al., 2009) |
| NP | 375 | E375G interacts with T-cell escape mutation at 384 (Rimmelzwaan et al., 2004; Gong et al., 2013) |
| NP | 384 | R384G and R384K are T-cell escape mutations (Voeten et al., 2000; Rimmelzwaan et al., 2004) |
| NP | 422 | K422R is a T-cell escape mutation (Boon et al., 2004) |
| HA | 138 | Contacts antigenic-site residues defined by the experiments of Caton et al. (1982) |
| HA | 189 | Contacts antigenic-site residues defined by the experiments of Caton et al. (1982) |
| HA | 225 | An antigenic site residue defined by the experiments of Caton et al. (1982) |

For lactamases, Salverda et al. (2010) report 18 sites at which mutations known to affect resistance are observed in clinical isolates. The ExpCM identify 9 sites of selection; 6 of these 9 sites are among the 18 known sites of resistance mutations (Table 3). There are 263 residues in the mature lactamase protein, so we can reject the possibility that the identified sites are not associated with resistance mutations (*P* = 10^−6^, Fisher’s exact test). So for lactamase, the ExpCM mostly identify sites that have been independently shown to affect drug resistance.

NP is under immune selection to escape T cells (Voeten et al., 2000; Machkovech et al., 2015) and probably also antibodies (Carragher et al., 2008; Laidlaw et al., 2013). The ExpCM identify 10 sites of selection. I searched the literature and found reports that 8 of these 10 sites are relevant to immune escape (Table 3). So for NP, the ExpCM mostly identify sites that have been independently shown to affect immunogenicity.

HA is under immune selection to escape antibodies. Caton et al. (1982) used antibodies to map escape mutations in H1 HA. A reasonable definition of the antigenic portion of HA is the set of sites identified by Caton et al. (1982) plus any sites in three-dimensional contact with these sites (a contact is defined as a *C*_*α*_ – *C*_*α*_ distance ≤ 6^°^*A* in PDB 1RVX). Using this definition, 86 of the 509 sites in the HA ectodomain are in the antigenic portion of the molecule. The ExpCM identify 3 sites of selection, all of which are in the antigenic portion of HA. We can reject the possibility that these identified sites are not associated with the antigenic portion of the molecule (*P* = 0.005, Fisher’s exact test). So for HA, the ExpCM identify sites that have been independently shown to affect immunogenicity.

Overall, these results show that sites of selection identified by ExpCM are indeed the locations of substitutions that alter evolutionarily relevant phenotypes. For a concrete illustration of sites of adaptive substitutions that are identified by ExpCM but not by a *dN*/*dS* method, Figure 5 shows the results of the ExpCM analysis of the five example sites in lactamase discussed in the Introduction and Figure 1. Three of these five sites experience substitutions that affect resistance, but a *dN*/*dS* method fails to flag any of them as under diversifying selection (*dN*/*dS* > 1) since it doesn’t account for site-specific constraints (Figure 1). Figure 5 shows that ExpCM correctly identify all three resistance sites as under diversifying or differential selection, while finding that the non-resistance sites are evolving as expected. Visual inspection of the two figures provides an intuitive explanation of why accounting for site-specific amino-acid preferences makes ExpCM so much more powerful at identifying sites of selection to alter evolutionarily relevant phenotypes.

**Figure 5:**
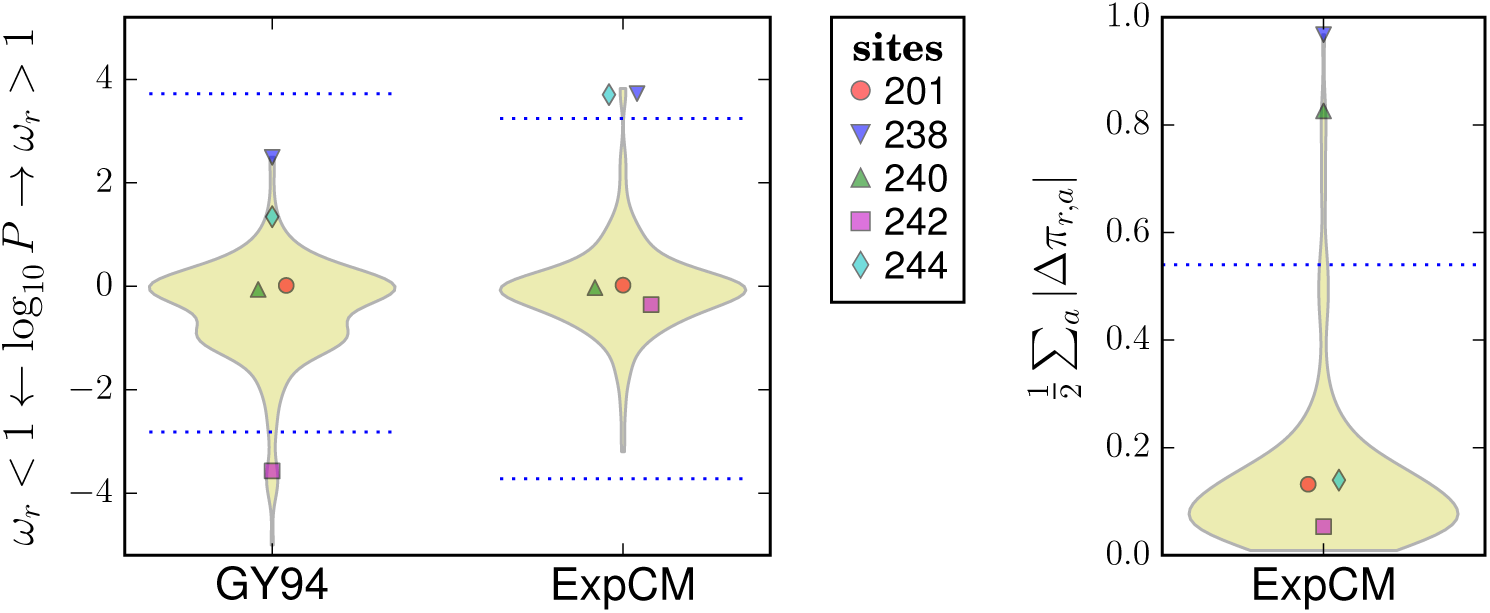
The experimentally informed models (ExpCM) correctly identify the three lactamase sites in Figure 1 that contribute to drug resistance. Figure 1 showed five sites in lactamase, three of which (238, 240, and 244) experience substitutions that contribute to drug resistance. However, a *dN*/*dS* analysis (GY94) fails to identify any of these sites as under diversifying selection (*dN*/*dS* > 1) at a FDR of 0.05 for testing all sites (dotted blue lines). In contrast, ExpCM correctly determine that the three resistance sites are under diversifying (238 and 244) or differential (238 and 240) selection, and that the two nonresistance sites (201 and 242) are evolving as expected. ExpCM outperform the *dN*/*dS* method because they implement a null model that accounts for the site-specific amino-acid preferences shown in Figure 1; for instance, this null model is not surprised that site 242 remains fixed at the highly preferred amino-acid R, but does find it noteworthy that site 240 substitutes to K multiple times even though that is not a particularly preferred amino acid.

## Discussion

I have described an approach that uses experimentally informed models to identify sites of biologically interesting selection in protein-coding genes. This approach asks the following question: *Is a site evolving differently in nature than expected from constraints measured in the lab?* In contrast, traditional *dN*/*dS* methods simply ask: *Is a site evolving non-neutrally?* The former question is sometimes more informative than the latter. It is by now abundantly clear that most protein residues are under some type of constraint, so finding that a site evolves non-neutrally is often unsurprising. Instead, we want to identify sites of substitutions that have altered evolutionarily relevant phenotypes. As demonstrated here, experimentally informed models have much greater power to identify such sites. The improvement is remarkable: while a *dN*/*dS* method fails to find any sites of adaptive evolution in the genes examined, experimentally informed models identify 22 sites of diversifying or differential selection, most of which fix mutations that have been independently shown to affect drug resistance or immunogenicity.

What accounts for the improve power of the experimentally informed site-specific models? As vividly illustrated by the deep mutational scanning studies that provide the data used here (Figure 2, Supplementary figure 3, Supplementary figure 4, and Supplementary figure 5), there is vast variation in the constraints on sites within a protein. Therefore, the significance that we should ascribe to a substitution depends on where it occurs: several changes at an unconstrained site may be unremarkable, but a single substitution away from a preferred amino acid at a constrained site probably reflects some powerful selective force. Whereas *dN*/*dS* methods treat all substitutions equally, the models used here evaluate the significance of each substitution in the context of the experimentally measured amino-acid preferences of the site at which it occurs.

Does this reliance on experimental measurements make the approach less objective? At first glance, the fact that *dN*/*dS* methods are uncontaminated by messy experiments feels reassuring. In contrast, experimentally informed models are dependent on all the subjective decisions associated with experimental design and interpretation. But in truth, experimentally informed models simply make explicit something that is already true: we define positive selection with respect to a null model for evolution in the absence of this selection. At least for the genes examined here, sites of known adaptive mutations are better identified by leveraging imperfect experiments that capture many of the constraints on natural evolution than by objectively testing the implausible null hypothesis that every site is evolving neutrally.

An assumption of experimentally informed site-specific models is that amino-acid preferences are conserved among the homologs under analysis. At first glance this assumption seems tenuous – epistasis can shift the effects of mutations as a gene evolves (Lunzer et al., 2010; Bridgham et al., 2009; Gong et al., 2013). But it is rare for epistatic shifts to be large enough to undermine the advantage of site-specific models: this fact is demonstrated by direct experiments (Doud et al., 2015; Risso et al., 2015; Ashenberg et al., 2013), the observation that parallel viral lineages tend to substitute to the same preferred amino acids at each site (Zanini et al., 2015), and the empirical superiority of site-specific models in fitting phylogenies of diverged homologs (Table 2; Bloom, 2014b; Doud et al., 2015). Therefore, epistasis does not subvert the basic advantage of a model informed by site-specific amino-acid preferences.

Of course, experimentally informed site-specific models require measurement of amino-acid preferences. However, advances in deep mutational scanning will make this requirement less and less of an impediment (Fowler and Fields, 2014; Boucher et al., 2014). In a fitting twist, one of the pioneers of deep mutational scanning (Fowler and Fields, 2014) was also the first to sequence a gene from influenza (Fields et al., 1981; Fields, 2016). At the time, sequencing the homologous gene from thousands of other viral strains must have seemed unimaginable – a few decades later, for this study I had to subsample the ≫10^5^ publicly available influenza sequences down to a manageable number. The core techniques of deep mutational scanning – sequencing and gene/genome engineering – are improving at a similar pace, so coming years will see measurement of the amino-acid preferences of many more genes.

Another possibility is to use non-experimental strategies to inform site-specific models like the one here. One strategy is to predict site-specific constraints from higher-level properties such as solvent accessibility (Meyer and Wilke, 2013; Shahmoradi et al., 2014; Meyer and Wilke, 2015) or via molecular simulation (Fornasari et al., 2002; Kleinman et al., 2010; Arenas et al., 2015; Echave et al., 2015). It remains unclear whether such non-experimental strategies can predict site-specific amino-acid preferences with sufficient accuracy to inform substitution models that can match the ExpCM used here. Another strategy is to infer preferences from naturally occurring sequences (Rodrigue et al., 2010; Rodrigue and Lartillot, 2014; Tamuri et al., 2012, 2014; Hopf et al., 2015). If these sequences can be partitioned into distinct clades, then preferences inferred from one clade might be used to identify selection acting only in another clade. But direct measurement via deep mutational scanning may well prove the best solution: after all, biology is full of properties that are challenging to predict or infer, but are now routinely measured in high-throughput.

Overall, I have described a new approach that leverages high-throughput experimental data to identify sites of selection in protein-coding genes. This approach clearly outperforms a standard implementation of the widely used *dN*/*dS* strategy, however there remains room for improvement. The utility of the *dN*/*dS* strategy has been enhanced by innovations that have made it possible to do things like test for selection only along certain branches (Murrell et al., 2012; Yang and Dos Reis, 2011), utilize Bayesian approaches to share information across sites (Nielsen and Yang, 1998; Yang et al., 2005; Huelsenbeck et al., 2006; Murrell et al., 2013), and more rapidly perform the computational analyses (Delport et al., 2010; Murrell et al., 2013). Most of these innovations could also be used in combination with the experimentally informed models described here. Methodological improvements of this sort, coupled with growing amounts of deep mutational scanning data, could make experimentally informed models an increasingly powerful tool to identify genotypic changes that have altered phenotypes of interest.

## Materials and Methods

## The phydms software package

The algorithms described in this paper are implemented in the `phydms` software package, which is freely available at https://github.com/jbloom/phydms. The `phydms` package is written in `Python`, and uses `cython` to interface with and extend `Bio`++ (http://biopp.univ-montp2.fr/; Dutheil et al., 2006; Guéguen et al., 2013) for the core likelihood calculations. Special thanks are due to Laurent Guéguen and Julien Dutheil for generously making the cutting-edge version of `Bio`++ available and providing assistance in its use. The `phydms` software also uses `dms_tools` (https://github.com/jbloom/dms_tools; Bloom, 2015) and `weblogo` (http://weblogo.threeplusone.com/; Crooks et al., 2004) for processing and visualizing the data and results.

## Computer code and data availability

The data and commands to perform the analyses and simulations in this paper are included as Supplementary Files with this preprint on `bioRxiv`. Within each archive, there is an explanatory `iPython` notebook that performs the analyses. The `ZIP` archives also contain the preferences, alignments, and full `phydms` output. Below is a brief description of the steps used to assemble the preferences and alignments.

## Amino-acid preferences for the four proteins

For NP, the preferences were taken directly from those provided by Doud et al. (2015), using the average of the measurements for the two NP variants. For HA, the preferences were taken directly from those provided by Thyagarajan and Bloom (2014). For lactamase, Stiffler et al. (2015) provide their data as “relative fitness” scores, which are the log_10_ of the enrichment ratios. I used the scores for the selections on 2.5 mg/ml of ampicillin (the highest concentration), averaging the scores for the two experimental trials. Following the definition of Bloom (2015) of the preferences as the normalized enrichment ratios, the preferences *π*_*r*_,_*a*_ are calculated from the relative fitness scores *S*_*r*_,_*a*_ so that *π*_*r*_,_*a*_ ∝ max (10^*S*_*r*,*a*_^, 10^−4^) and 1 = Σ_*a*_*π*_*r*_,_*a*_. For Gal4, Kitzman et al. (2015) provide their data as “effect scores”, which are the log_2_ of the enrichment ratios. The preferences are calculated from the effect scores *E*_*r*_,_*a*_ so that *π*_*r*_,_*a*_ ∝ max (2^*E*_*r*,*a*_^, 2 × 10^−4^) and 1 = Σ_*a*_ *π*_*r*_,_*a*_. A few of the effect scores are missing from Kitzman et al. (2015), so these scores are set to the average score for all mutations for which scores are provided. The formulas to convert the lactamase and Gal4 scores to preferences include the max operators to avoid estimating preferences of zero; the minimal allowable values specified by the second argument to these operators are my guess of the lowest frequency that would have been reliably observed in each experiment.

## Alignments

For NP, the sequence alignment was constructed by extracting all post-1950 full-length NPs in the Influenza Virus Resource (Bao et al., 2008) that are descended in purely human lineages from the 1918 virus (this is H1N1 from 1950-1957 and 1977-2008, H2N2 from 1957-1968, and H3N2 from 1968-2015), and then retaining just two sequences per-subtype per-year to yield a manageable alignment. The rationale for using only post-1950 sequences is that most influenza sequences isolated before that date were passaged extensively in the lab prior to sequencing. For HA, the alignment was constructed by extracting all post-1950 sequences in the human seasonal H1N1 lineage (this is H1N1 from 1950-1957 and 1977-2008), and then retaining just four sequences per year to yield a manageable alignment. For lactamase, the alignment consists of the TEM and SHV lactamases used in Bloom (2014b). For Gal4, a set of homologs was obtained by performing a `tblastn` search of the Gal4 DNA-binding domain used Kitzman et al. (2015) against `wgs` (limiting by `saccharomyceta` (`taxid:716545`)) and `chromosomes` for hits with *E* ≤ 0.01, and then retaining only sequences that aligned to the Gal4 DNA-binding domain with ≥ 70% protein identity and ≤ 5% gaps. For all genes, alignments were made pairwise to the sequence used for the deep mutational scanning with `EMBOSS needle` (Rice et al., 2000), and any sites were purged if they were gapped in that reference sequence.

## Sequence numbering

In the figures and tables, the residues in NP are numbered sequentially beginning with one at the N-terminal methionine. The residues in HA are numbered using the H3 numbering scheme (this is the one used in PDB 4HMG), and the site-specific selection analysis is performed only for the residues in HA ectodomain (these are the residues present in PDB 4HMG). The residues in lactamase are numbered using the scheme of Ambler et al. (1991). The residues in Gal4 are numbered using the same scheme as in Kitzman et al. (2015).

## Acknowledgments

Tremendous thanks to Laurent Guéguen and Julien Dutheil for developing the `Bio`++ libraries, generously making the cutting-edge version of this software freely available, and providing assistance in its use. Thanks to Erick Matsen for helpful suggestions about software and coding. Thanks to Sergei Kosakovsky-Pond and an anonymous reviewer for helpful comments on the first draft of the manuscript. This work was supported by the NIGMS of the NIH under grant R01GM102198.

---

**Supplementary figure 1:**
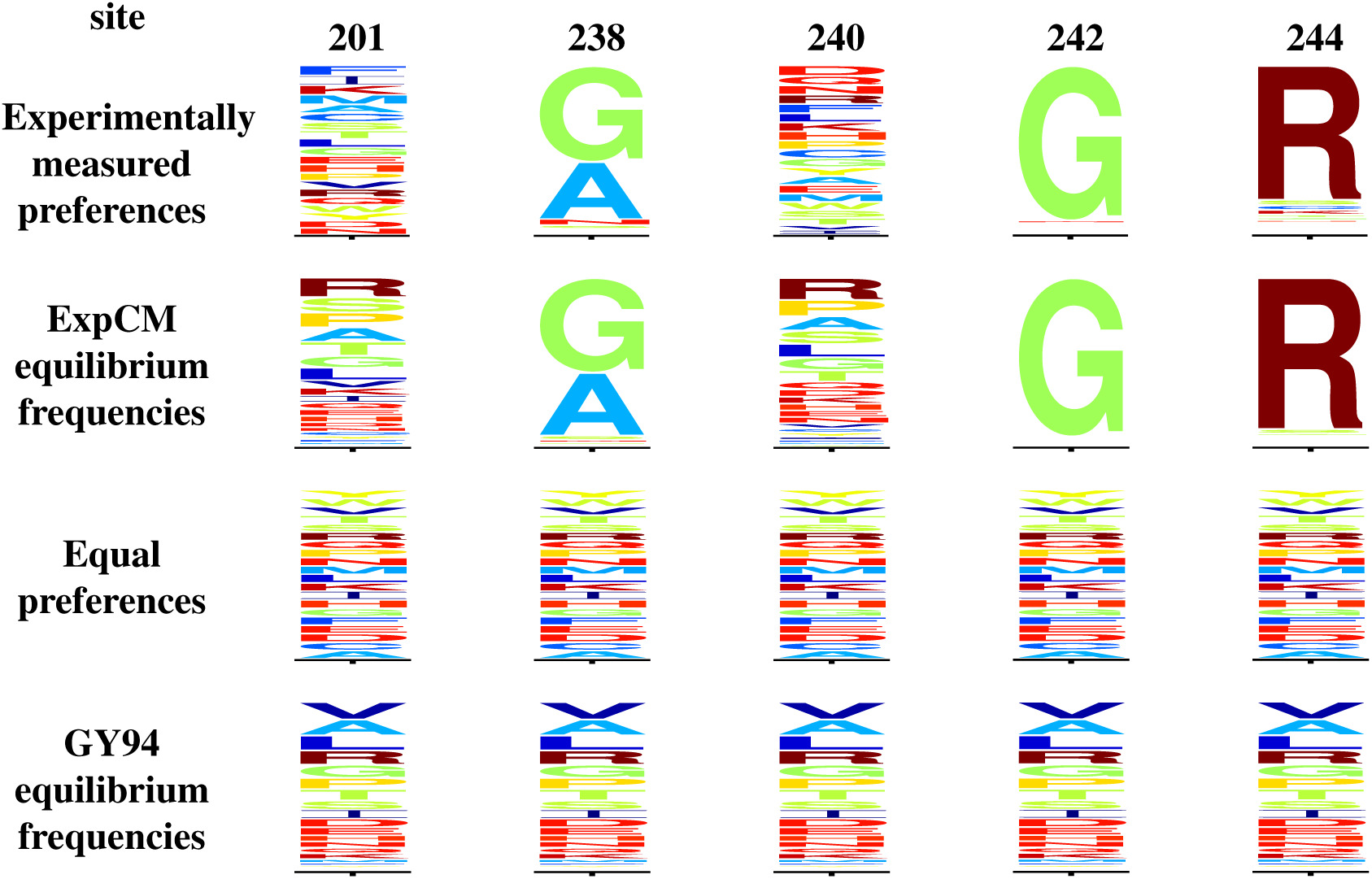
Clarification of subtleties in the relationship between amino-acid preferences and substitution model equilibrium frequencies. Figure 1 shows the experimentally measured amino-acid preferences and the equilibrium frequencies of the GY94 model. The equilibrium frequencies of the experimentally informed codon models (ExpCM) are given by Equation 4, and are similar but not identical to the preferences: the ExpCM equilibrium frequencies are also influenced by the unequal number of codons per amino acid, nucleotide mutation biases, and the stringency parameter *β*. The equilibrium frequencies of the GY94 model already account for the codon/mutation factors. To clarify these distinctions, this figure shows the preferences and equilibrium frequencies of the ExpCM model, and the “all-equal” amino-acid preferences that would lead to the equilibrium frequencies of the GY94 model if the nucleotide frequency parameters in that model are construed as representing mutation-level rather than selection-level processes. Note that the logo plots show the amino-acid frequencies implied by the equilibrium codon frequencies (i.e. the sum of the frequencies of all encoding codons for each amino acid).

**Supplementary figure 2:**
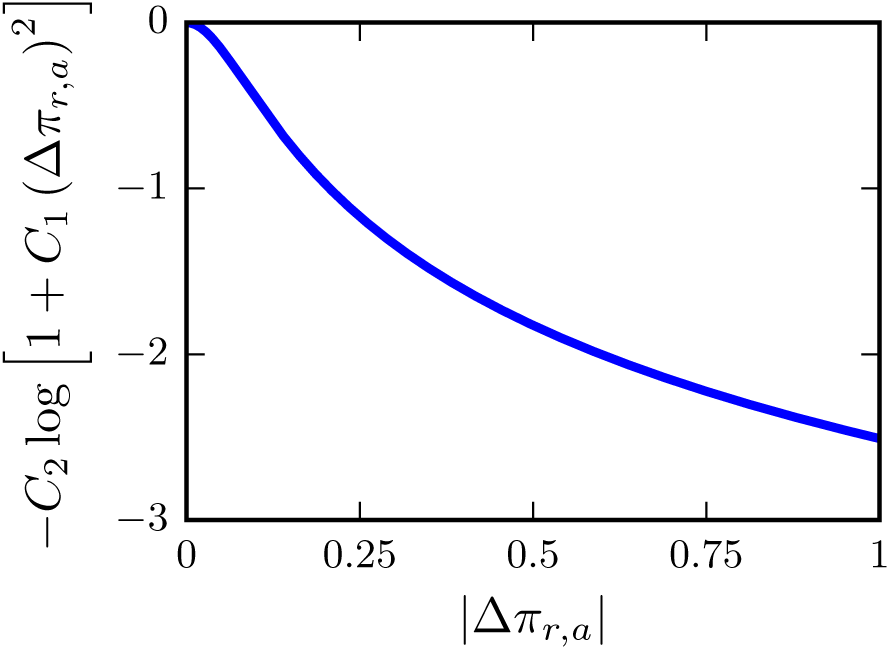
Graphical illustration of the equation used to regularize the Δ*π*_*r*,*a*_ values when inferring differential selection. The log of the regularization defined by Equation 7 is a sum of terms like this taken over all differential preferences at a site. This regularization has the property that the marginal cost of shifting Δ*π*_*r*_,_*a*_ away from zero is initially steep but then flattens somewhat as Δ*π*_*r*_,_*a*_ becomes large. This corresponds to the intuition that most sites will be evolving as expected (and so have Δ*π*_*r*_,_*a*_ ~ 0), but a few sites might be under strong differential selection. This plot uses *C*_1_ = 150 and *C*_2_ = 0.5.

**Supplementary figure 3:**
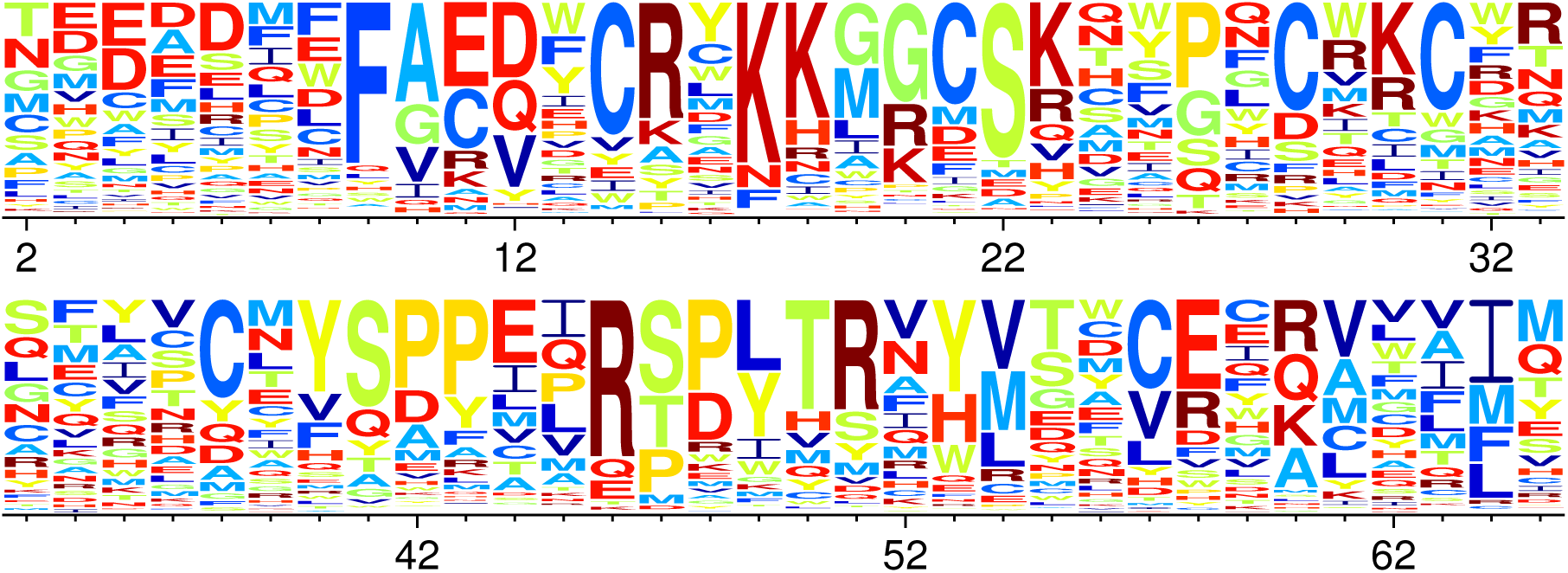
Site-specific amino-acid preferences for Gal4. Shown are the preferences experimentally measured by Kitzman et al. (2015) for the DNA-binding domain of yeast Gal4, re-scaled by the stringency parameter *β* = 0.82 from Table 2.

**Supplementary figure 4:**
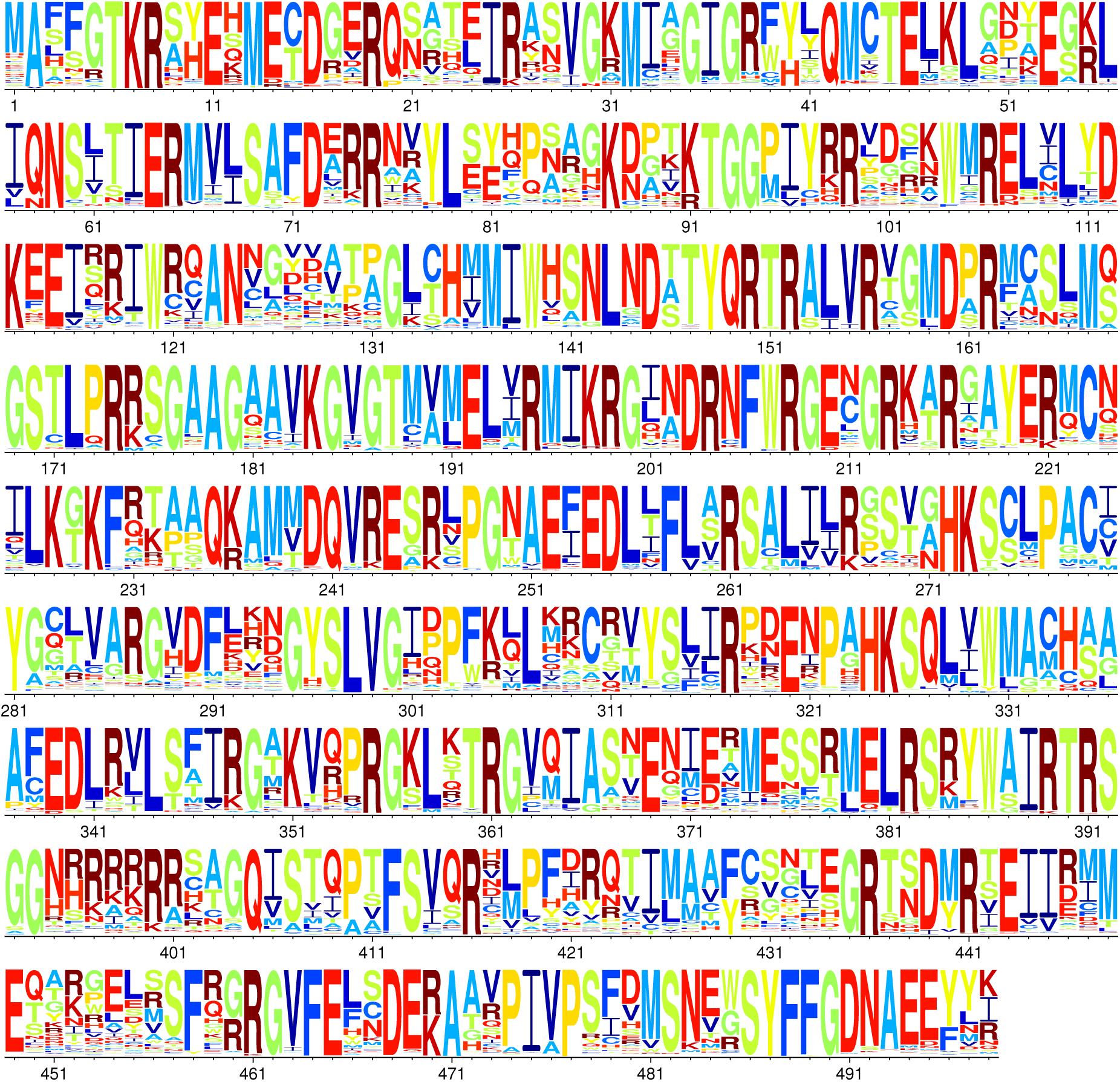
Site-specific amino-acid preferences for influenza NP. Shown are the preferences experimentally reported in Doud et al. (2015) for the average of the measurements on the A/PR/8/1934 and A/Aichi/2/1968 strains, re-scaled by the stringency parameter *β* = 2.43 from Table 2.

**Supplementary figure 5:**
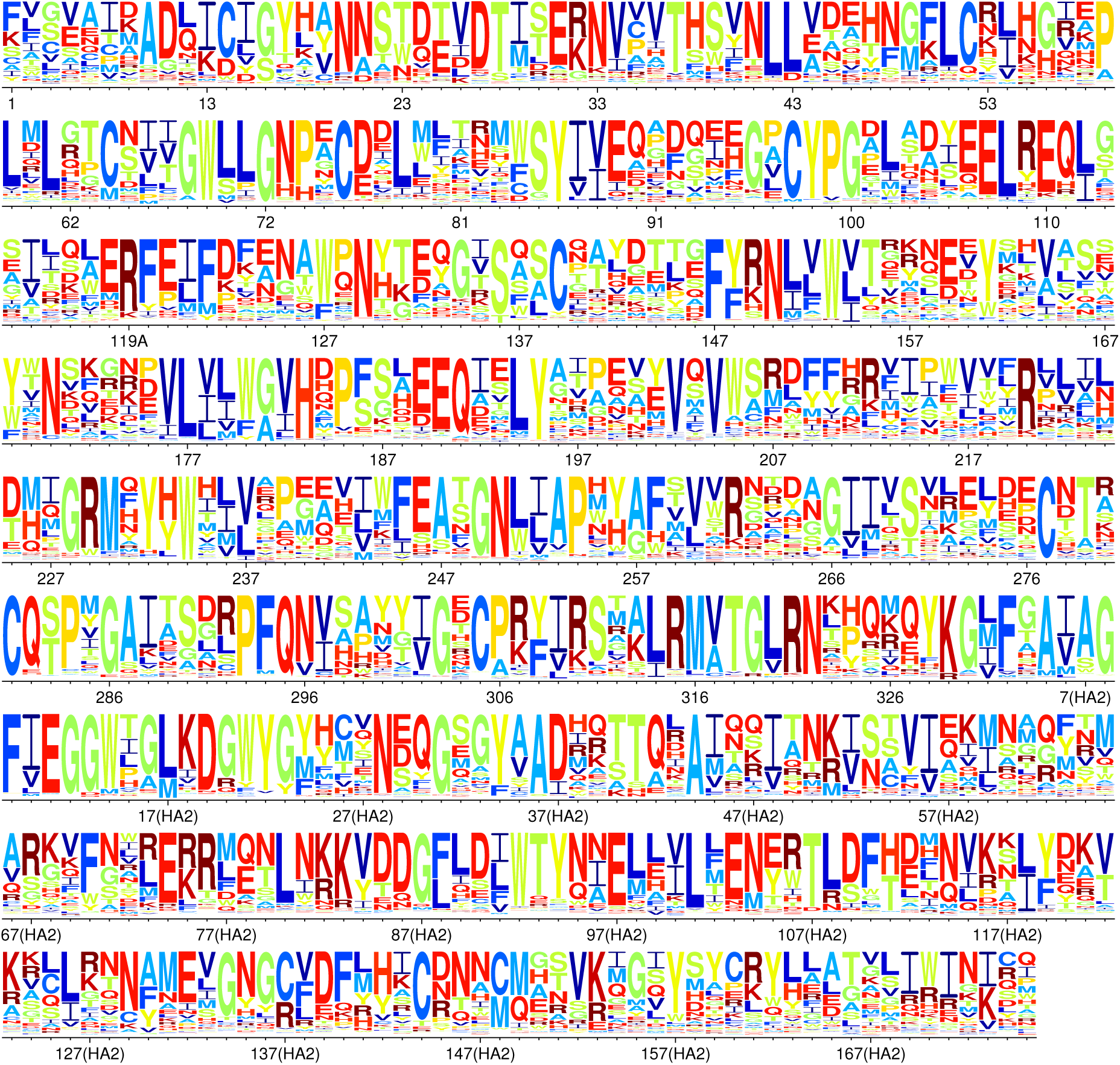
Site-specific amino-acid preferences for HA. Shown are the preferences experimentally measured by Thyagarajan and Bloom (2014) for influenza HA (A/WSN/1933, H1N1 strain), re-scaled by the stringency parameter *β* = 1.61 from Table 2. The residues are numbered according to the H3 numbering scheme (the one used in PDB 4HMG), and data are only shown for sites in the HA ectodomain (residues present in the crystal structure in PDB 4HMG).

**Supplementary figure 6:**
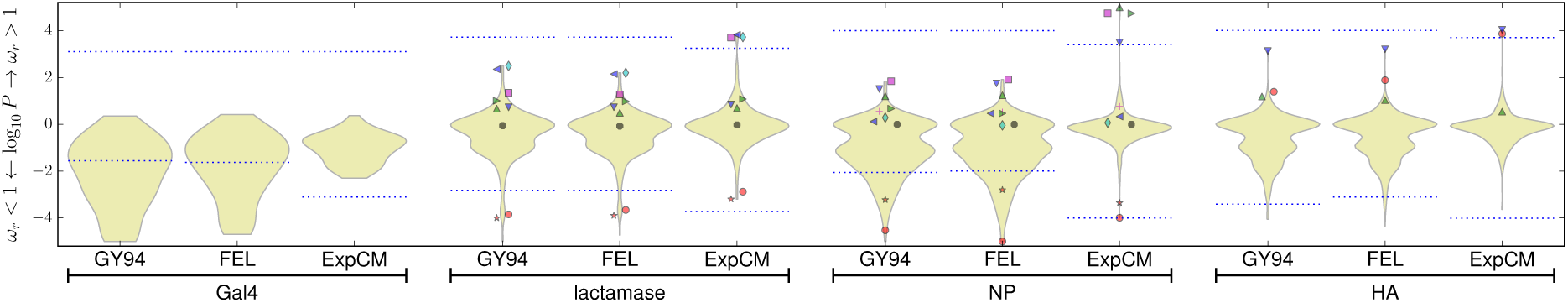
The results of the *dN*/*dS* analysis are qualitatively similar when using `HyPhy` rather than `phydms`. This figure shows the same data as that in Figure 3A, but also includes the results of a *dN*/*dS* analysis using the fixed effects likelihood (FEL) method implemented in `HyPhy` (Pond et al., 2005). The results are not identical to the `phydms` GY94 results because the `HyPhy` implementation differs slightly from the `phydms` implementation: `HyPhy` performs the *dN*/*dS* analysis using the substitution model of Muse and Gaut (1994) rather than GY94, and infers a neighbor-joining tree with a nucleotide substitution model rather than a maximum-likelihood tree using a codon model. Nonetheless, the results of the `HyPhy` FEL analysis are highly similar to those of the `phydms` GY94 analysis, both in terms of the overall distribution of results and in terms of the values for the specific indicated sites. The point markers represent the same sites as in Figure 3.

**Supplementary figure 7:**
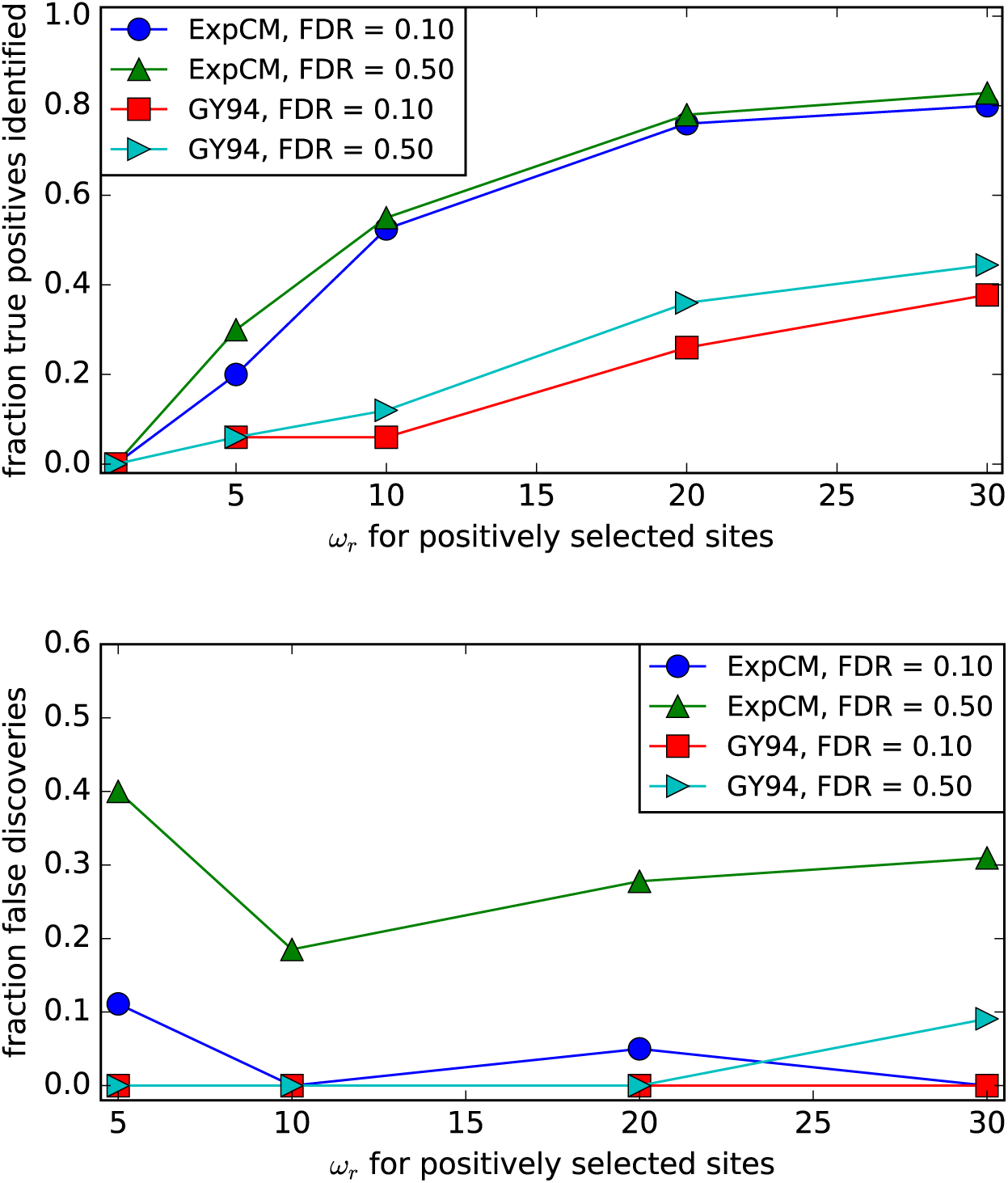
Simulations validate the statistical approach used to identify diversifying selection. Using the actual ExpCM parameters for NP in Table 2 except fixing *ω* = 1 for all sites except for those selected to be simulated under diversifying selection, I used `pyvolve` (Spielman and Wilke, 2015b) to simulate 10 alignments along tree inferred from the actual NP sequences. For each simulation, I randomly selected 5 sites to place under diversifying selection, with *ω*_*r*_ values ranging from 1 (no diversifying selection) to 30 (very strong diversifying selection). I then analyzed the data using `phydms` in the same way that the actual data were analyzed. Sites were called as being under significant diversifying selection using the false discovery rates (FDRs) indicated in the figure. The top panel shows that ExpCM greatly outperformed the FEL-like GY94 method at identifying true positives. The bottom panel shows that the Benjamini and Hochberg (1995) procedure effectively controls the fraction of false discoveries among the sites called as being under diversifying selection using ExpCM. The Benjamini and Hochberg (1995) procedure may be slightly too conservative for ExpCM (for every value of *ω*_*r*_ the actual rate of false discoveries is slightly below the FDR), but the differences seem modest.

**Supplementary figure 8:**
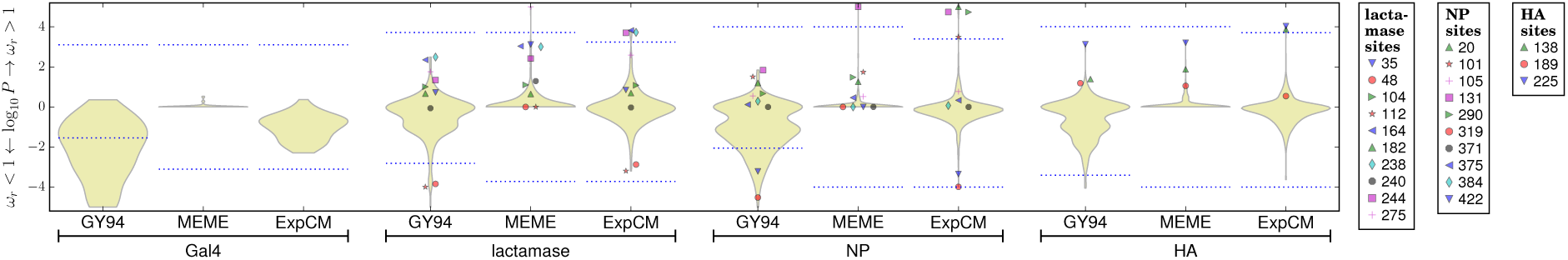
This figure is same as Figure 3A but also includes an analysis with `MEME` (Murrell et al., 2012) as implemented in `HyPhy` (Pond et al., 2005). `MEME` reports the P-value that a site has *dN*/*dS* > 1 on at least some branches of the tree. As can be seen from this figure, `MEME` is somewhat more powerful than the GY94-based FEL approach, presumably because some sites are only under episodic diversifying selection. While the GY94-based FEL approach identifies no sites of diversifying selection, `MEME` identifies one site of diversifying selection in lactamase and one site in NP. However, `MEME` still identifies fewer sites for all genes than the ExpCM.

**Supplementary figure 9:**
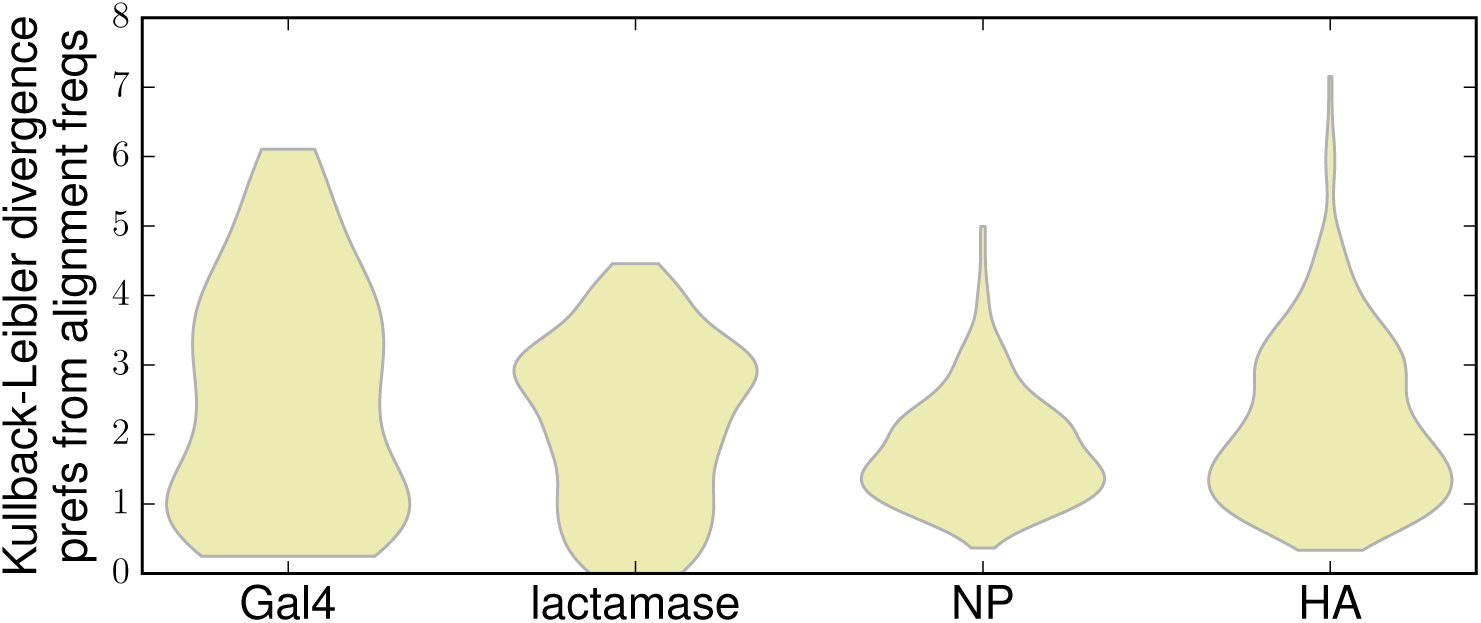
This figure shows the distribution over sites of the Kullback-Leibler divergence of the experimentally measured amino-acid preferences from the alignment frequencies. Note that the Kullback-Leibler divergence does *not* take phylogeny into account, and so will be confounded the incomplete sampling of potentially tolerated amino acids by natural evolution. The distribution of per-site Kullback-Leibler divergences shown here lacks the biologically sensible features of the differential selection computed in a phylogenetic framework and shown in Figure 3B. For instance, Gal4 has many sites with very high Kullback-Leibler divergence even though on biological grounds we expect it to be evolving mostly in the absence of positive selection. In contrast, lactamase and NP tend to have lower Kullback-Leibler divergence even though we know that they evolve under selection for adaptive mutations that confer drug resistance or immune escape. The biologically unreasonable distribution of Kullback-Leibler divergences shown in this plot are probably due to the failure of the Kullback-Leibler divergence to account for phylogeny, which may in turn make the results highly sensitive to uneven phylogenetic sampling and differences in the total sequence divergence spanned by the alignments (see Table 1). The Kullback-Leibler divergence was computed using logarithms taken to the base two.

**Supplementary figure 10:**
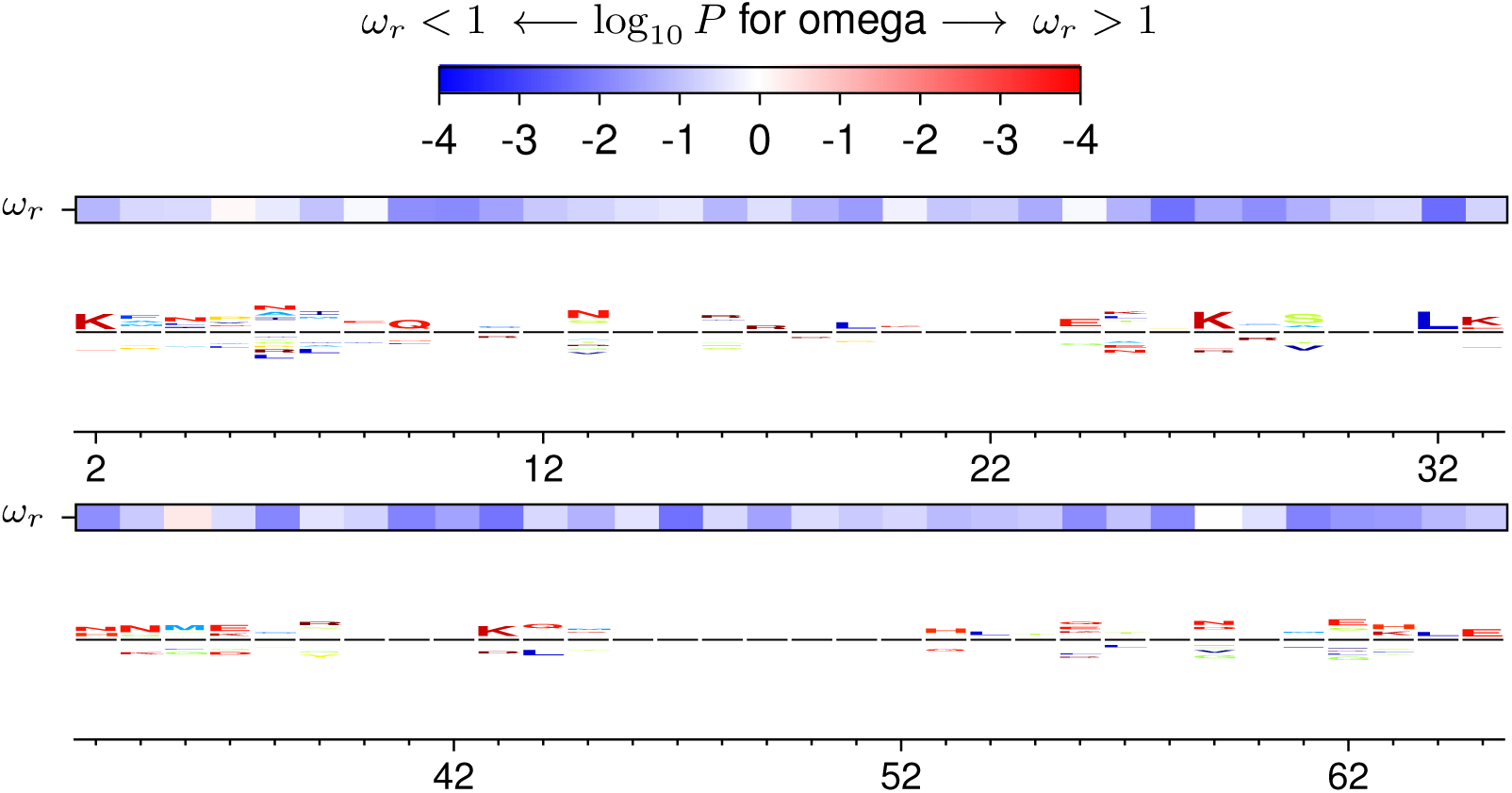
Site-specific selection on Gal4 inferred with the experimentally informed models.

**Supplementary figure 11:**
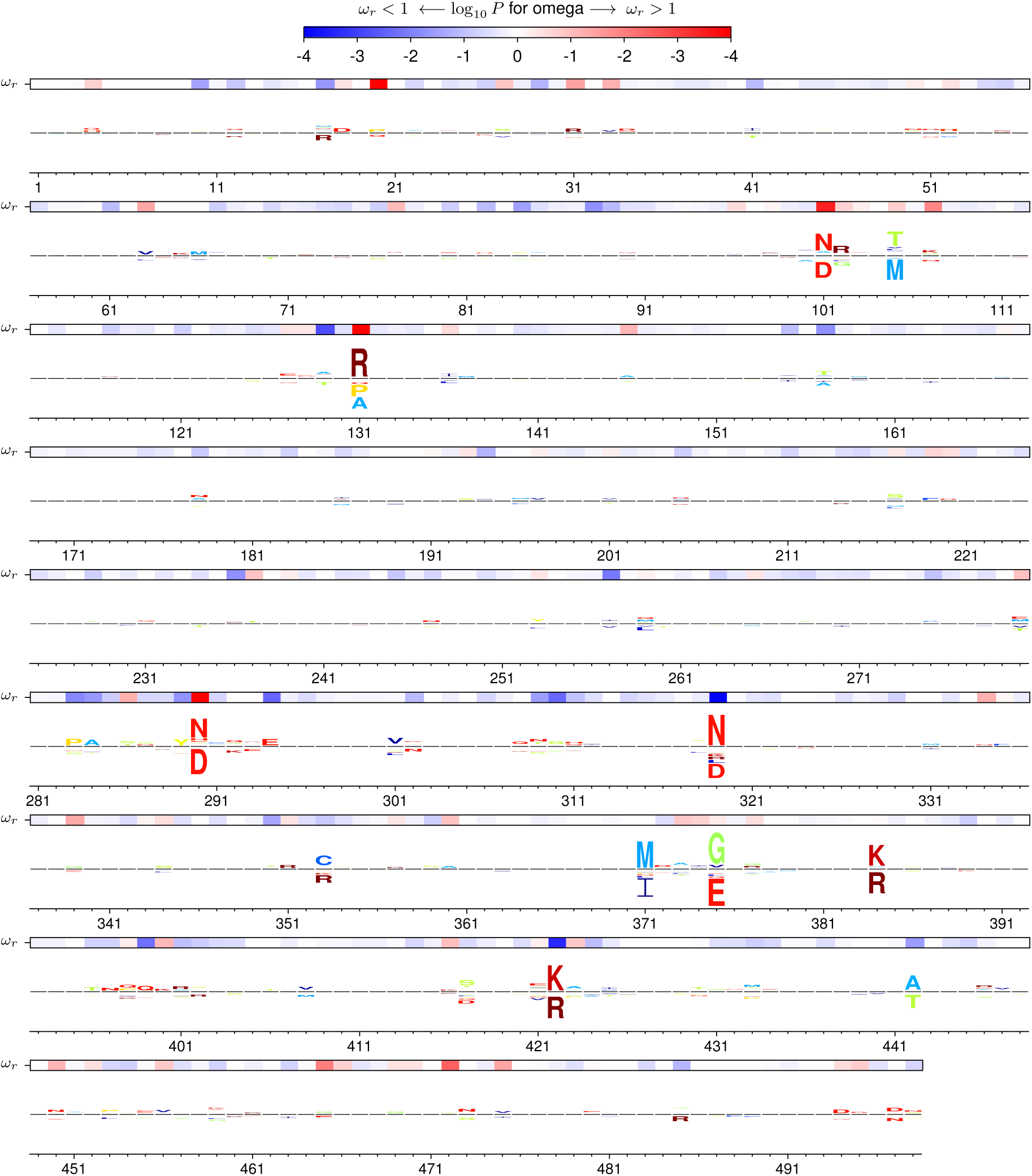
Site-specific selection on NP inferred with the experimentally informed models.

**Supplementary figure 12:**
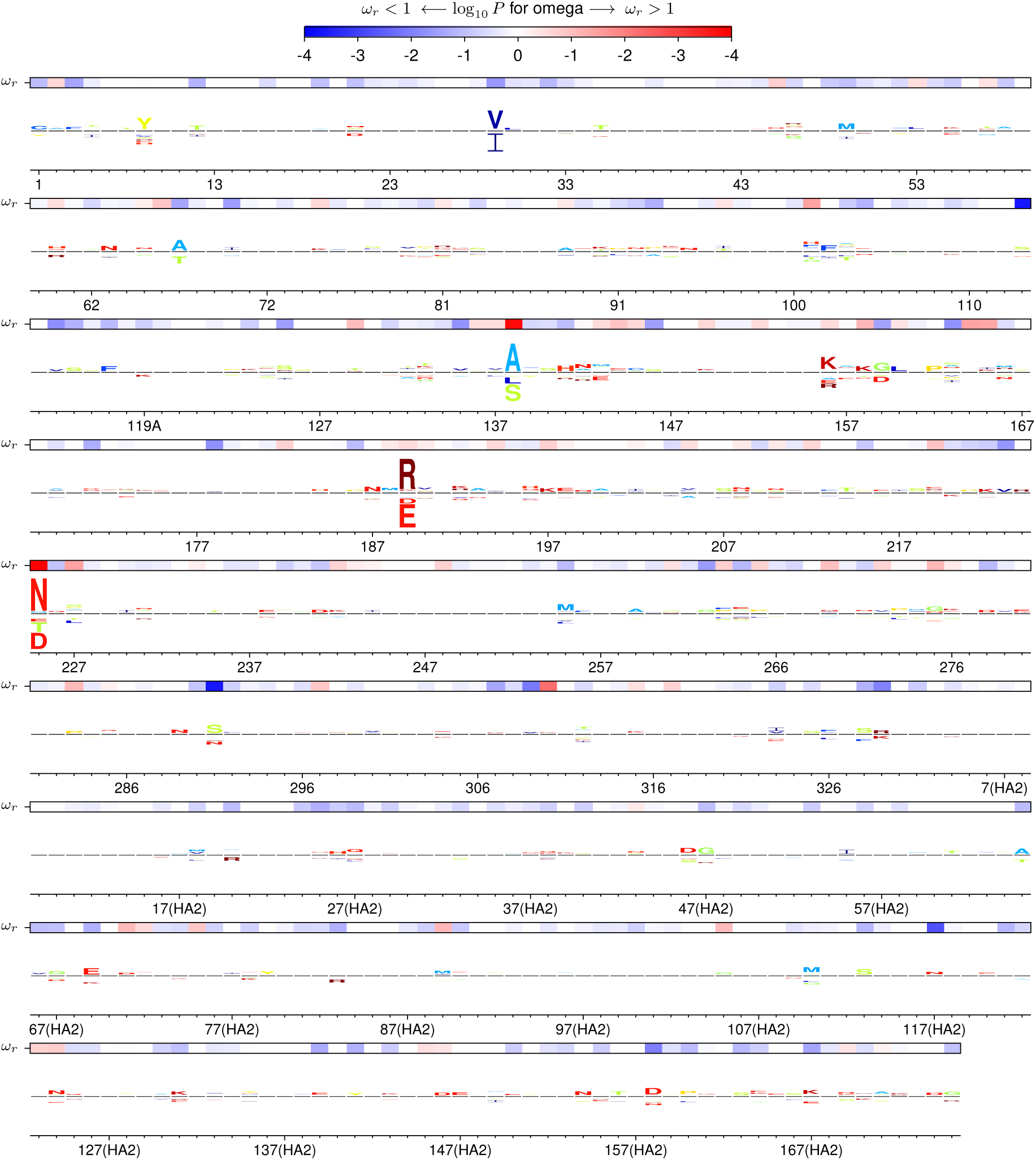
Site-specific selection on HA inferred with the experimentally informed codon models. Residues are numbered as in Supplementary figure 5.

## References

Alexander J, Bilsel P, del Guercio MF, et al. (11 co-authors). 2010. Identification of broad binding class I HLA supertype epitopes to provide universal coverage of influenza A virus. Human immunology. 71:468–474.

Ambler R, Coulson A, Frére JM, Ghuysen JM, Joris B, Forsman M, Levesque R, Tiraby G, Waley S. 1991. A standard numbering scheme for the class A beta-lactamases. Biochemical Journal. 276:269.

Arenas M, Sánchez-Cobos A, Bastolla U. 2015. Maximum likelihood phylogenetic inference with selection on protein folding stability. Molecular Biology and Evolution. 32:2195–2207.

Ashenberg O, Gong LI, Bloom JD. 2013. Mutational effects on stability are largely conserved during protein evolution. Proc. Natl. Acad. Sci. USA. 110:21071–21076.

Bao Y, Bolotov P, Dernovoy D, Kiryutin B, Zaslavsky L, Tatusova T, Ostell J, Lipman D. 2008. The Influenza Virus Resource at the National Center for Biotechnology Information. J. Virol. 82:596–601.

Benjamini Y, Hochberg Y. 1995. Controlling the false discovery rate: a practical and powerful approach to multiple testing. Journal of the Royal Statistical Society. Series B. Methodological. 57:289–300.

Berkhoff E, Geelhoed-Mieras M, Fouchier R, Osterhaus A, Rimmelzwaan G. 2007. Assessment of the extent of variation in influenza A virus cytotoxic T-lymphocyte epitopes by using virus-specific CD8+ T-cell clones. Journal of General Virology. 88:530–535.

Bloom JD. 2014a. An experimentally determined evolutionary model dramatically improves phylogenetic fit. Molecular Biology and Evolution. 31:1956–1978.

Bloom JD. 2014b. An experimentally informed evolutionary model improves phylogenetic fit to divergent lactamase homologs. Molecular Biology and Evolution. 31:2753–2769.

Bloom JD. 2015. Software for the analysis and visualization of deep mutational scanning data. BMC Bioinformatics. 16:168.

Boon AC, de Mutsert G, van Baarle D, Smith DJ, Lapedes AS, Fouchier RA, Sintnicolaas K, Osterhaus AD, Rimmelzwaan GF. 2004. Recognition of homo-and heterosubtypic variants of influenza A viruses by human CD8+ T lymphocytes. The Journal of Immunology. 172:2453–2460.

Boucher JI, Cote P, Flynn J, Jiang L, Laban A, Mishra P, Roscoe BP, Bolon DN. 2014. Viewing protein fitness landscapes through a next-gen lens. Genetics. 198:461–471.

Bridgham JT, Ortlund EA, Thornton JW. 2009. An epistatic ratchet constrains the direction of glucocorticoid receptor evolution. Nature. 461:515–519.

Carragher DM, Kaminski DA, Moquin A, Hartson L, Randall TD. 2008. A novel role for non-neutralizing antibodies against nucleoprotein in facilitating resistance to influenza virus. The Journal of Immunology. 181:4168–4176.

Caton AJ, Brownlee GG, Yewdell JW, Gerhard W. 1982. The antigenic structure of the influenza virus A/PR/8/34 hemagglutinin (H1 subtype). Cell. 31:417–427.

Crooks GE, Hon G, Chandonia JM, Brenner SE. 2004. Weblogo: a sequence logo generator. Genome Research. 14:1188–1190.

Delport W, Poon AF, Frost SD, Pond SLK. 2010. Datamonkey 2010: a suite of phylogenetic analysis tools for evolutionary biology. Bioinformatics. 26:2455–2457.

Doud MB, Ashenberg O, Bloom J. 2015. Site-specific amino-acid preferences are mostly conserved in two closely related protein homologs. Molecular Biology and Evolution. 32:2944–2960.

Du Bois S, Marriott M, Amyes S. 1995. TEM-and SHV-derived extended-spectrum β-lactamases: relationship between selection, structure and function. Journal of Antimicrobial Chemotherapy. 35:7–22.

Dutheil J, Gaillard S, Bazin E, Glémin S, Ranwez V, Galtier N, Belkhir K. 2006. Bio++: a set of C++ libraries for sequence analysis, phylogenetics, molecular evolution and population genetics. BMC Bioinformatics. 7:188.

Echave J, Jackson EL, Wilke CO. 2015. Relationship between protein thermodynamic constraints and variation of evolutionary rates among sites. Physical biology. 12:025002.

Echave J, Spielman S, Wilke C. 2016. Causes of evolutionary rate variation among protein sites. Nature reviews. Genetics. 17:109–121.

Felsenstein J. 1981. Evolutionary trees from DNA sequences: a maximum likelihood approach. Journal of Molecular Evolution. 17:368–376.

Fields S. 2016. Count em. https://genestogenomes.org/count-em/.

Fields S, Winter G, Brownlee GG. 1981. Structure of the neuraminidase gene in human influenza virus A/PR/8/34. Nature. 290:213–217.

Fornasari MS, Parisi G, Echave J. 2002. Site-specific amino acid replacement matrices from structurally constrained protein evolution simulations. Mol. Biol. Evol. 19:352–356.

Fowler DM, Fields S. 2014. Deep mutational scanning: a new style of protein science. Nature Methods. 11:801–807.

Goldman N, Yang Z. 1994. A codon-based model of nucleotide substitution probabilities for protein-coding DNA sequences. Mol. Biol. Evol. 11:725–736.

Gong LI, Suchard MA, Bloom JD. 2013. Stability-mediated epistasis constrains the evolution of an influenza protein. eLife. 2:e00631.

Guéguen L, Gaillard S, Boussau B, et al. (11 co-authors). 2013. Bio++: Efficient extensible libraries and tools for computational molecular evolution. Molecular Biology and Evolution. 30:1745–1750.

Guindon S, Rodrigo AG, Dyer KA, Huelsenbeck JP. 2004. Modeling the site-specific variation of selection patterns along lineages. Proceedings of the National Academy of Sciences. 101:12957–12962.

Guo HH, Choe J, Loeb LA. 2004. Protein tolerance to random amino acid change. Proc. Natl. Acad. Sci. USA. 101:9205–9210.

Halpern AL, Bruno WJ. 1998. Evolutionary distances for protein-coding sequences: modeling site-specific residue frequencies. Mol. Biol. Evol. 15:910–917.

Hasegawa M, Kishino H, Yano Ta. 1985. Dating of the human-ape splitting by a molecular clock of mitochondrial DNA. Journal of Molecular Evolution. 22:160–174.

Hopf TA, Ingraham JB, Poelwijk FJ, Springer M, Sander C, Marks DS. 2015. Quantification of the effect of mutations using a global probability model of natural sequence variation. arXiv preprint arXiv:1510.04612. .

Huelsenbeck JP, Jain S, Frost SW, Pond SLK. 2006. A dirichlet process model for detecting positive selection in protein-coding dna sequences. Proceedings of the National Academy of Sciences. 103:6263–6268. .

Johnston M. 1987. A model fungal gene regulatory mechanism: the GAL genes of saccharomyces cerevisiae. Microbiological reviews. 51:458.

Kitzman JO, Starita LM, Lo RS, Fields S, Shendure J. 2015. Massively parallel single-amino-acid mutagenesis. Nature Methods. 12:203–206.

Kleinman CL, Rodrigue N, Lartillot N, Philippe H. 2010. Statistical potentials for improved structurally constrained evolutionary models. Mol. Biol. Evol. 27:1546–1560.

Laidlaw BJ, Decman V, Ali M, et al. (11 co-authors). 2013. Cooperativity between CD8+ T cells, nonneutralizing antibodies, and alveolar macrophages is important for heterosubtypic influenza virus immunity. PLoS Pathogens. 9:e1003207.

Lunzer M, Golding GB, Dean AM. 2010. Pervasive cryptic epistasis in molecular evolution. PLoS Genetics. 6:e1001162.

Machkovech HM, Bedford T, Suchard MA, Bloom JD. 2015. Positive selection in CD8+ T-cell epitopes of influenza virus nucleoprotein revealed by a comparative analysis of human and swine viral lineages. Journal of Virology. 89:11275–11283.

Massingham T, Goldman N. 2005. Detecting amino acid sites under positive selection and purifying selection. Genetics. 169:1753–1762.

McCandlish DM, Stoltzfus A. 2014. Modeling evolution using the probability of fixation: History and implications. The Quarterly Review of Biology. 89:225–252.

Meyer AG, Wilke CO. 2013. Integrating sequence variation and protein structure to identify sites under selection. Molecular biology and evolution. 30:36–44.

Meyer AG, Wilke CO. 2015. The utility of protein structure as a predictor of site-wise dn/ds varies widely among hiv-1 proteins. Journal of The Royal Society Interface. 12:20150579.

Miyoshi-Akiyama T, Yamashiro T, Mai LQ, Narahara K, Miyamoto A, Shinagawa S, Mori S, Kitajima H, Kirikae T. 2012. Discrimination of influenza A subtype by antibodies recognizing host-specific amino acids in the viral nucleoprotein. Influenza and Other Respiratory Viruses. 6:434–441.

Murrell B, Moola S, Mabona A, Weighill T, Sheward D, Kosakovsky PS, Scheffler K. 2013. FUBAR: a fast, unconstrained bayesian approximation for inferring selection. Molecular Biology and Evolution. 30:1196–1205.

Murrell B, Wertheim J, Moola S, Weighill T, Scheffler K, Kosakovsky PS. 2012. Detecting individual sites subject to episodic diversifying selection. PLoS Genetics. 8:e1002764.

Muse SV, Gaut BS. 1994. A likelihood approach for comparing synonymous and nonsynonymous nucleotide substitution rates, with application to the chloroplast genome. Molecular Biology and Evolution. 11:715–724.

Nielsen R, Yang Z. 1998. Likelihood models for detecting positively selected amino acid sites and applications to the HIV-1 envelope gene. Genetics. 148:929–936.

Pond SK, Muse SV. 2005. Site-to-site variation of synonymous substitution rates. Molecular Biology and Evolution. 22:2375–2385.

Pond SL, Frost SD, Muse SV. 2005. HyPhy: hypothesis testing using phylogenies. Bioinformatics. 21:676–679.

Pond SLK, Frost SD. 2005. Not so different after all: a comparison of methods for detecting amino acid sites under selection. Molecular Biology and Evolution. 22:1208–1222.

Posada D, Buckley TR. 2004. Model selection and model averaging in phylogenetics: advantages of Akaike information criterion and Bayesian approaches over likelihood ratio tests. Systematic Biology. 53:793–808.

Rice P, Longden I, Bleasby A. 2000. EMBOSS: the European molecular biology open software suite. Trends in Genetics. 16:276–277.

Rimmelzwaan G, Berkhoff E, Nieuwkoop N, Fouchier R, Osterhaus A. 2004. Functional compensation of a detrimental amino acid substitution in a cytotoxic-T-lymphocyte epitope of influenza A viruses by co-mutations. Journal of Virology. 78:8946–8949.

Risso V, Manssour-Triedo F, Delgado-Delgado A, et al. (11 co-authors). 2015. Mutational studies on resurrected ancestral proteins reveal conservation of site-specific amino acid preferences throughout evolutionary history. Molecular Biology and Evolution. 32:440–455.

Rodrigue N, Lartillot N. 2014. Site-heterogeneous mutation-selection models within the PhyloBayes-MPI package. Bioinformatics. 30:1020–1021.

Rodrigue N, Philippe H, Lartillot N. 2010. Mutation-selection models of coding sequence evolution with site-heterogeneous amino acid fitness profiles. Proceedings of the National Academy of Sciences. 107:4629–4634.

Salverda ML, De Visser JAG, Barlow M. 2010. Natural evolution of TEM-1 *β*-lactamase: experimental reconstruction and clinical relevance. FEMS Microbiology Reviews. 34:1015–1036.

Shahmoradi A, Sydykova DK, Spielman SJ, Jackson EL, Dawson ET, Meyer AG, Wilke CO. 2014. Predicting evolutionary site variability from structure in viral proteins: buriedness, packing, flexibility, and design. Journal of Molecular Evolution. 79:130–142.

Spielman S, Wilke C. 2015a. The relationship between *dn*/*ds* and scaled selection coefficients. Molecular biology and evolution. 32:1097–1108.

Spielman SJ, Wilke CO. 2015b. Pyvolve: a flexible Python module for simulating sequences along phylogenies. PloS One. 10:e0139047.

Stiffler MA, Hekstra DR, Ranganathan R. 2015. Evolvability as a function of purifying selection in TEM-1 *β*-lactamase. Cell. 160:882–892.

Suzuki Y. 2004. New methods for detecting positive selection at single amino acid sites. Journal of molecular evolution. 59:11–19.

Suzuki Y, Gojobori T. 1999. A method for detecting positive selection at single amino acid sites. Molecular Biology and Evolution. 16:1315–1328.

Tamuri AU, dos Reis M, Goldstein RA. 2012. Estimating the distribution of selection coefficients from phylogenetic data using sitewise mutation-selection models. Genetics. 190:1101–1115.

Tamuri AU, Goldman N, dos Reis M. 2014. A penalized likelihood method for estimating the distribution of selection coefficients from phylogenetic data. Genetics. pp. genetics–114.

Thorne JL, Choi SC, Yu J, Higgs PG, Kishino H. 2007. Population genetics without intraspecific data. Molecular Biology and Evolution. 24:1667–1677.

Thyagarajan B, Bloom JD. 2014. The inherent mutational tolerance and antigenic evolvability of influenza hemagglutinin. eLife. 3:e03300.

Traven A, Jelicic B, Sopta M. 2006. Yeast Gal4: a transcriptional paradigm revisited. EMBO reports. 7:496–499.

Varich N, Kaverin N. 2004. Antigenically relevant amino acid positions as revealed by reactions of monoclonal antibodies with the nucleoproteins of closely related influenza A virus strains. Archives of virology. 149:2271–2276.

Varich NL, Kochergin-Nikitsky KS, Usachev EV, Usacheva OV, Prilipov AG, Webster RG, Kaverin NV. 2009. Location of antigenic sites recognized by monoclonal antibodies in the influenza A virus nucleoprotein molecule. Journal of General Virology. 90:1730–1733.

Varich NL, Sadykova GK, Prilipov AG, Kochergin-Nikitsky KS, Kushch AA, Masalova OV, Klimova RR, Gitelman AK, Kaverin NV. 2011. Antibody-binding epitope differences in the nucleoprotein of avian and mammalian influenza A viruses. Viral immunology. 24:101–107.

Voeten J, Bestebroer T, Nieuwkoop N, Fouchier R, Osterhaus A, Rimmelzwaan G. 2000. Antigenic drift in the influenza A virus (H3N2) nucleoprotein and escape from recognition by cytotoxic T lymphocytes. Journal of Virology. 74:6800–6807.

Yang Z. 2007. PAML 4: phylogenetic analysis by maximum likelihood. Molecular Biology and Evolution. 24:1586–1591.

Yang Z, Dos Reis M. 2011. Statistical properties of the branch-site test of positive selection. Molecular biology and evolution. 28:1217–1228.

Yang Z, Nielsen R. 2002. Codon-substitution models for detecting molecular adaptation at individual sites along specific lineages. Molecular biology and evolution. 19:908–917.

Yang Z, Nielsen R. 2008. Mutation-selection models of codon substitution and their use to estimate selective strengths on codon usage. Molecular biology and evolution. 25:568–579.

Yang Z, Nielsen R, Goldman N, Pedersen AMK. 2000. Codon-substitution models for heterogeneous selection pressure at amino acid sites. Genetics. 155:431–449.

Yang Z, Wong WS, Nielsen R. 2005. Bayes empirical bayes inference of amino acid sites under positive selection. Molecular Biology and Evolution. 22:1107–1118.

Yewdell J, Webster R, Gerhard W. 1979. Antigenic variation in three distinct determinants of an influenza type A haemagglutinin molecule. Nature. 279:246–248.

Zanini F, Brodin J, Thebo L, Lanz C, Bratt G, Albert J, Neher RA. 2015. Population genomics of intrapatient HIV-1 evolution. eLife. p. e11282.

Zuckerkandl E, Pauling L. 1965. Evolutionary divergence and convergence in proteins. In: Evolving genes and proteins. New York, NY: Academic Press, pp. 97–166.

